# Enhanced Recombination Among SARS-CoV-2 Omicron Variants Contributes to Viral Immune Escape

**DOI:** 10.1101/2022.08.23.504936

**Authors:** Rishad Shiraz, Shashank Tripathi

## Abstract

SARS-CoV-2 virus evolution occurs as a result of antigenic drift and shift. Although antigenic drift has been extensively studied, antigenic shift, which for SARS-CoV-2 occurs through genetic recombination, has been examined scarcely. To gain a better understanding of the emergence and prevalence of recombinant SARS-CoV-2 lineages through time and space, we analyzed SARS-CoV-2 genome sequences from public databases. Our study revealed an extraordinary increase in the emergence of SARS-CoV-2 recombinant lineages during the Omicron wave, particularly in Northern America and Europe. This phenomenon was independent of sequencing density or genetic diversity of circulating SARS-CoV-2 strains. In SARS-CoV-2 genomes, recombination breakpoints were found to be more concentrated in the 3’ UTR followed by ORF1a. Additionally, we noted enrichment of certain amino acids in the spike protein of recombinant lineages, which have been reported to confer immune escape from neutralizing antibodies, increase ACE2 receptor binding, and enhance viral transmission in some cases. Overall, we report an important and timely observation of accelerated recombination in the currently circulating Omicron variants and explore their potential contribution to viral fitness, particularly immune escape.

## INTRODUCTION

RNA viruses constitute the majority of emerging and re-emerging human pathogens. These viruses are known to accumulate genetic mutations at a higher rate, compared to other infectious agents with DNA genomes (1). This is primarily due to the error-prone action of viral RNA-dependent RNA polymerase (RdRP) and the lack of viral proofreading enzymes. Such mutations are known as ‘Antigenic Drift’ which is gradual, incremental and provides the genetic diversity essential for viral fitness. These mutations contribute to viral zoonosis, immune escape, enhanced transmission, altered tropism and pathogenesis (2–4). SARS-CoV-2 has a ~30 kb long ssRNA genome which is also replicated by an error-prone RdRP (NSP12) introducing 8×10^-4^ nucleotide substitution/site/year(5). Although SARS-CoV-2 encodes for an exonuclease enzyme (NSP14) with proofreading ability, it has shown remarkable antigenic drift to evade infection and vaccine-mediated immunity and enhance viral transmission during progressive waves by different variants of concern (VOCs) (6–12). Another way, by which sudden large-scale genetic changes appear in the RNA virus genomes is called ‘Antigenic Shift’, which happens either by genetic reassortment or by genetic recombination (13–15). Genetic reassortment is known to underlie the emergence of Influenza A virus pandemic strains, however, it is limited to viruses with a segmented genome (13, 16). Genetic recombination, on the other hand, can happen in both segmented as well as non-segmented viral genomes. It has been reported to contribute to viral adaptation in cases of Polio, HIV, and HCV (14, 17, 18).

Genetic recombination in the RNA virus genomes happens through molecular processes such as template switching and homologous recombination (19). It requires co-infection of the host with parent strains, which are usually co-circulating in the same location (20). Genetic recombination is a shared feature of Sarbecovirus evolution and is believed to have contributed to the emergence of SARS-CoV, MERS as well as SARS-CoV-2 (21, 22). Recombination events in the SARS-CoV-2 genome during COVID-19 pandemic have been examined before in specific contexts (15, 20, 23–25). The first recombinant lineage reported, named XA appeared first in the UK and continued to circulate for a limited time (20). Later, recombinant lineage XB sequences were reported in the USA though it emerged before XA, with substantial forward circulation (15). Recombinants between VOCs were reported later, such as XC parented by Alpha and Delta that emerged in Japan, although with limited forward circulation (23).

A comprehensive analysis of the current status of recombinant SARS-CoV-2 lineages, their evolutionary history, phylogenetic relationship and contribution to viral evolution during the COVID-19 pandemic has been lacking. To understand the prevalence and significance of genetic recombination events in the SARS-CoV-2 genome, we analysed the publicly available whole genome datasets, spanning the entire COVID-19 pandemic. We observed a striking escalation in the appearance of recombinant lineage during the Omicron wave, although the first major recombinant lineage appeared during the Alpha wave. Geographically, the majority of recombinant lineages emerged in Northern America and Northern European countries, especially in the UK, subsequently spreading to different parts of the world. Detailed analysis of the nucleotide sequences of recombinant lineages revealed the untranslated regions (UTRs), especially the 3’ UTR to be a recombination hotspot. Among coding regions, recombination breakpoints were most prevalent in ORF1a. At the protein level, we observed conserved specific amino acid changes in the NSP14 exonuclease of recombinant lineages parented by Omicron VOC, which may have a potential role in the enhanced recombination frequency. Interestingly there were multiple mutations enriched in the Spike protein of recombinant lineages, which have been reported to provide resistance against neutralizing antibodies, strengthen ACE2 receptor binding and enhance viral transmission. Overall, this study provides timely observation of escalation in the appearance of recombinant SARS-CoV-2 lineages during Omicron wave and provides detailed insight into the functional relevance of genetic changes acquired through recombination, especially in immune escape.

## RESULTS

### SARS-CoV2 Omicron Variant wave coincided with an extraordinary escalation in the emergence of recombinant lineages

To understand the role of recombination in SARS-CoV-2 evolution, we analysed 1,206,055 complete SARS-CoV-2 genome sequences deposited in the NCBI database and all the recombinant lineage sequences deposited in the GISAID database collected between December 2019 to July 2022 (26, 27). Although recombination is one of the important strategies utilised by RNA viruses, SARS-CoV2 unlike other coronaviruses showed modest recombination (24), with only three recombinant lineages reported in the first 2 years of the pandemic, up to November 2021 [FIG1A]. However, subsequently in the next seven months, between December 2021 to July 2022, 28 new recombinant lineages emerged, tallying the total number of recombinant lineages from 3 to 31. This was a more than 1 order of magnitude increase in the number of new recombinant lineages. Next, we checked the timeline of appearance and period of circulation of the recombinant lineages based on data available in the GISAID (26). We observed that the first recombinant lineage of SARS-CoV2 was XB which appeared in July 2020 and was prevalent till September 2021. During these 5-months, two more recombinant lineages emerged, XA in January 2021 and XC in August 2021. Both these lineages had a limited circulation period of around 2 months. The last XC lineage sequence was collected in October 2021, and for the next 2 months till mid-December 2021, no recombinant lineage sequences were detected. However subsequently, the number of recombinant lineages escalated rapidly. In December 2021, three new recombinant lineages XT, XF and XH emerged. While January of 2022 recorded the emergence of 11 new recombinant lineages namely XG, XAC, XD, XS, XJ, XE, XU, XN, XM, XAH and XV, February of 2022 recorded 9 new recombinant lineages namely XAB, XL, XK, XQ, XAA, XR, XAF, XAD and XAE, March of 2022 recorded 2 recombinant lineages namely XZ and XY, and remaining XAG recombinant lineage emerged in April 2022. This increased frequency of recombination events, coincided with the peak of the Omicron wave, especially between December 2021 to January 2022 [S1A]. Next challenge was to understand whether detection of enhanced SARS-CoV-2 recombination events was due to a natural increase in the emergence of recombinant lineages or was a by-product of increased worldwide SARS-CoV-2 sequencing. For this, we examined the percentage prevalence of recombinant lineages sequence submissions over the COVID-19 pandemic timeline [S1B]. For each of the 29 recombinant lineages other than XD and XT, when they recorded maximum percentage prevalence, more than 1 in every 1000 SARS-CoV2 genomes sequenced belonged to these recombinant lineages. In the cases of XD and XT, they reported a sequence detection frequency of more than 1 in every 2000 SARS-CoV2 genomes sequenced during their peak percentage prevalence. These frequencies are suggestive of recombinant lineage detections not being an associated consequence of increased sequence surveillance efforts, but indeed due to increased natural emergence of recombinant lineages. To understand the prevalence of specific recombinant lineages through the pandemic, we analysed the total number of recombinant sequences from different lineages (in GISAID) over time [FIG1B]. The XB recombinant lineage, first emerged in July 2020 (earlier than alpha variant-driven pandemic waves), and it peaked in July of 2021 constituting ~ 9% of all SARS-CoV2 sequences [S1B]. Another recombinant lineage with a higher prevalence was XE, which was still circulating in July 2022. To understand the relationship between the emergence of recombinant lineages with other significant non-recombinant lineages, we compared the timeline of their emergence and circulation during the COVID-19 pandemic. Results showed a clear regime change in December 2021, with the fall of delta variant lineages and the emergence of omicron variant lineages [FIG1C]. It was evident that the surge of the omicron variant perfectly coincided with the increased emergence of recombinant lineages [S1A]. Furthermore, we examined the total number of unique SARS-CoV-2 lineages present at any given time during the pandemic, to see whether increased recombination was linked to increased available genetic diversity [S1C]. We did not observe any increase in the number of lineages detected per day during the rapid upsurge in recombinant lineages. This suggests the sudden extraordinary upsurge in the emergence of recombinant lineages during the omicron wave of the pandemic is not due to the overall increase in genetic diversity of SARS-CoV-2 lineages.

**Figure 1:**
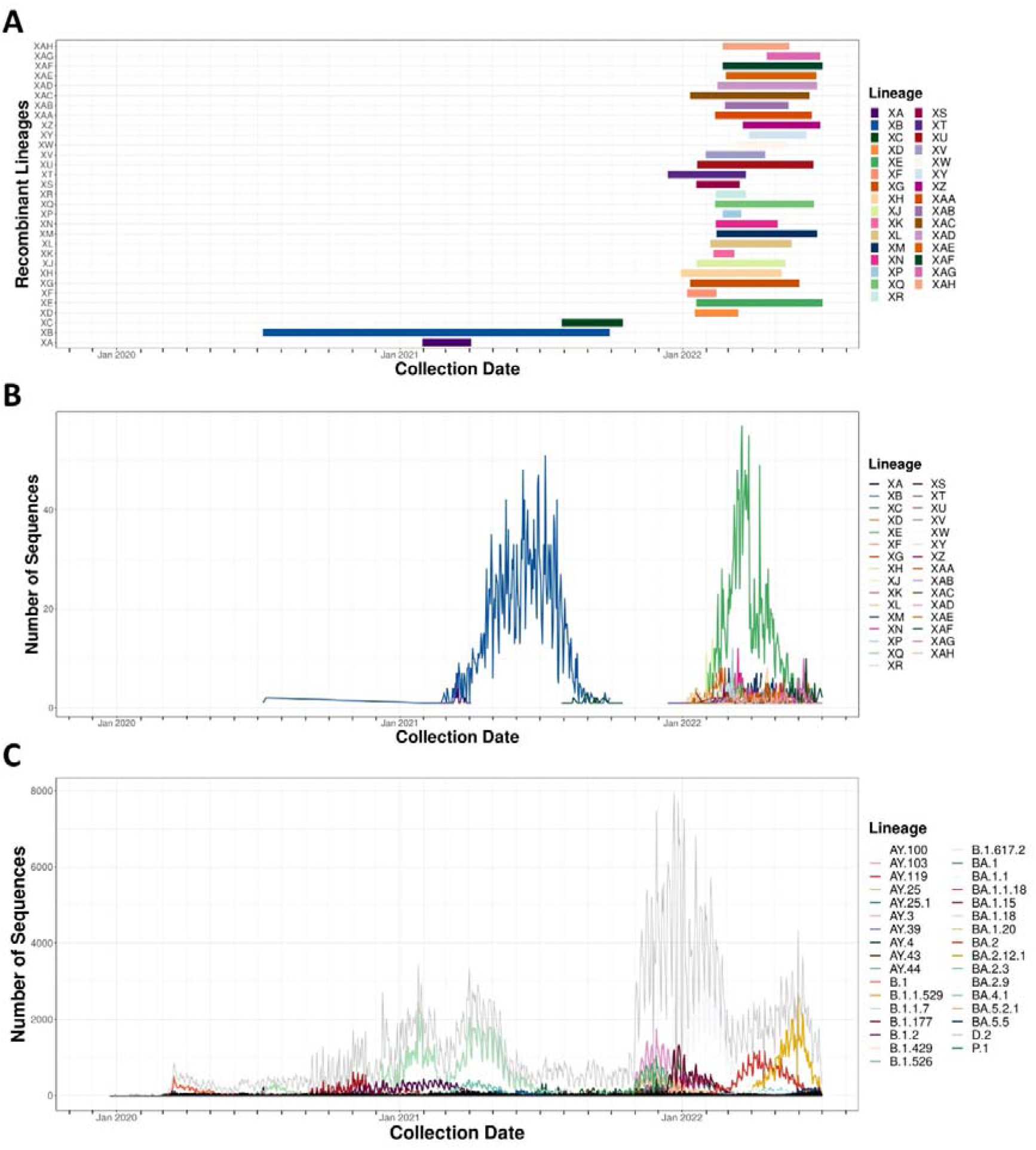
Timeline of SARS-CoV-2 recombinant lineage emergence and circulation. (A) Timeline of SARS-CoV2 recombinant lineages with Y axis showing recombinant lineages, X-axis showing the sequence collection date. Horizontal coloured bars indicate the period between the first and last detection of the corresponding colour-coded recombinant lineage. (B) Prevalence of SARS-CoV-2 recombinant lineages over time with Y axis showing the daily number of sequences reported, X-axis showing the sequence collection date having axis ticks indicating months. Each coloured line in the graph shows the daily number of sequences collected belonging to the corresponding recombinant lineage, with lineage colours the same as Fig1A. (C) Prevalence of all lineages of SARS-CoV-2 over time with Y axis showing a daily number of sequences reported, X-axis showing the sequence collection date having axis ticks indicating months. The Grey line in the graph represents the total number of sequences collected, while less prevalent lineages with less than 150 sequences collected on the day of its maximum peak are represented in black. Each coloured line other than black and grey in the graph represents major SARS-CoV-2 lineages with at least 150 of the lineage sequence collected on the day of its maximum peak. X axis of all three graphs ranges from 1^st^ December 2019 to 1^st^ July 2022 with axis ticks indicating months and are aligned making them temporally comparable to each other.

### Europe and North America have been the Hotspots for the emergence and spread of SARS-CoV-2 recombinant Lineages

Before the Omicron wave, recombinant lineages of SARS-CoV-2 were few and had limited geographical spread. To understand if the sudden increase in recombinant lineages during the Omicron wave was localized to specific regions, we analysed the geographic distribution of recombinant lineages compared to the cumulative spread of the SARS-CoV2 through the COVID-19 pandemic [FIG2A]. We observed that, although recombinant lineages spread across the world, they were more concentrated in Europe, North and Central American regions, followed by Asia, South America and least Australia and Africa. To test for possible geographic bias in recombinant lineage emergence, we analysed the percentage distribution of each recombinant lineage per country followed by marking the country of first detection [FIG2B]. As sequencing efforts of SARS-CoV2 genomes are geographically skewed with developed countries contributing more to the sequencing data (28), the first detection of recombinant lineage sequence in a country could be a mere consequence of higher sequencing efforts. But if a country recorded both first detection and the highest prevalence of a particular recombinant lineage, that country is identified as the country of emergence for that recombinant lineage. Out of the 28 recombinant lineages detected post-emergence of omicron variant, 16 lineages emerged in European countries as they were first detected and most prevalent in these countries. Out of these 16 recombinant lineages, a maximum number of lineages (7) namely XE, XF, XL, XN, XP, XQ and XR emerged in the UK, and 3 lineages XM, XAB and XAD emerged in Germany, and another 3 lineages XG, XH and XV emerged in Denmark, XJ emerged in Finland while XAH emerged from Slovenia. The XK recombinant lineage emerged in Belgium and was the only recombinant lineage that did not spread beyond the country of the first detection. Out of 28 recombinant lineages detected post-emergence of Omicron, 4 lineages namely XS, XY, XZ and XAA were first detected and remained most prevalent in the USA, and XAF emerged from Costa Rica representing Central America.

**Figure 2:**
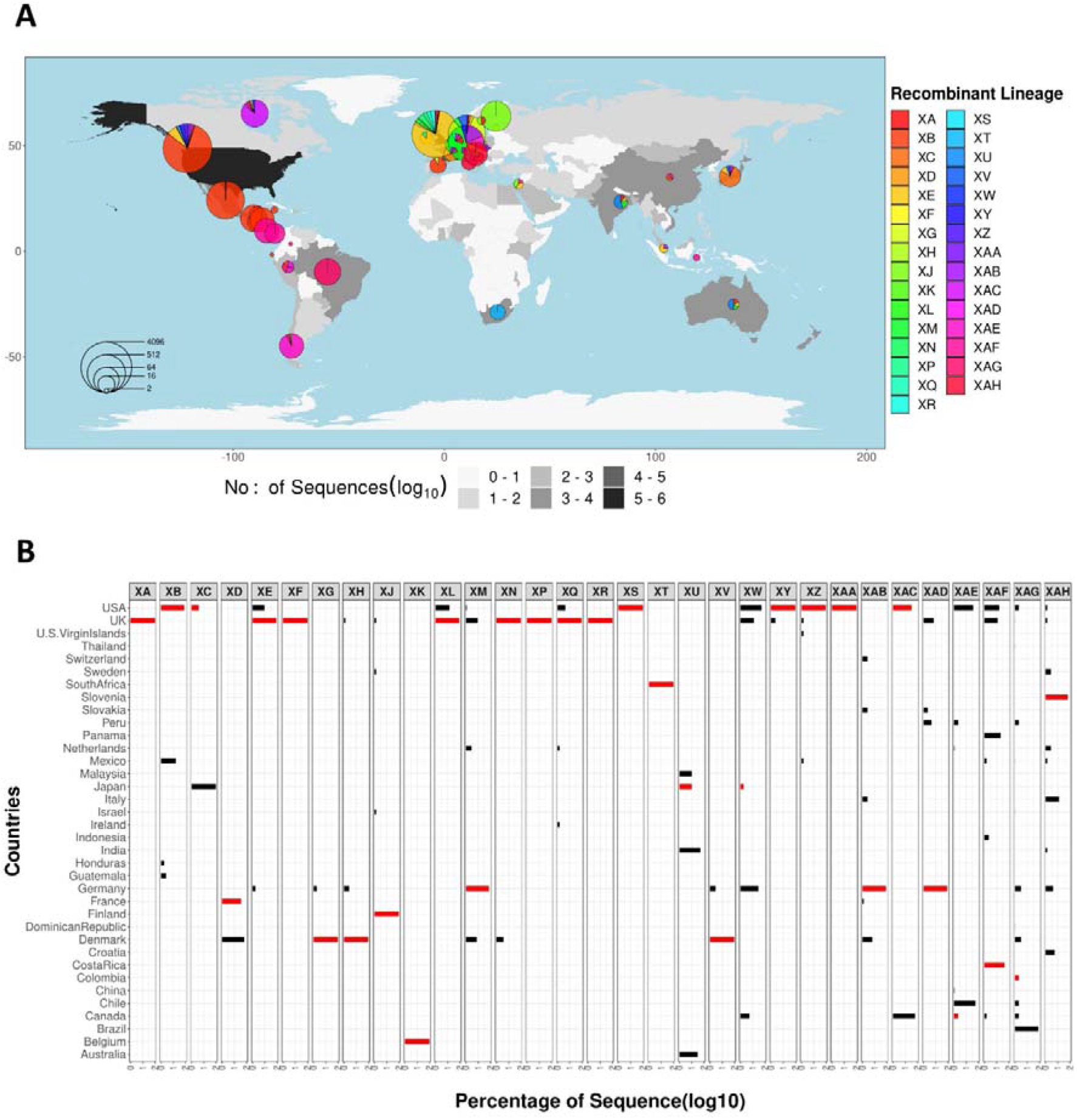
Global geographic spread, percentage geographic distribution and country of the first detection of SARS-CoV2 recombinant lineages. (A) Geographic distribution of SARS-CoV2 recombinant lineages with X and Y axis representing longitude and latitude respectively. The map fill colour of each country is a gradient of grey representing the number of SARS-CoV2 sequences collected and deposited onto the NCBI database from each country (in log10 scale) with dark grey representing more sequences (Legends positioned at the bottom). The pie chart in each country represents the distribution of recombinant lineage sequences collected from that country, with different colours representing different recombinant lineages (Legends positioned on the right side). The radius of the pie chart is proportional to the log2 number of all recombinant lineage sequences collected from that country(Legends positioned inside the map in the bottom left). (B) Facetted horizontal Bar graph representing percentage geographic distribution of each recombinant lineage, with lineage names corresponding to the facet mentioned on the top. For each facet, X-axis shows the percentage of that lineage sequence(in log10 scale) and the Y axis shows countries with at least 10% of any recombinant lineage sequence. Bars marked in red indicate the country from where the first sequence of that recombinant lineage was collected.

The XT lineage was exclusively detected in South Africa, representing the African continent. In cases of XD, XU, XW, XAC, XAE and XAG, although they were first detected in France, Japan, Japan, USA, Canada and Colombia respectively, they showed maximum prevalence in Denmark, India, USA, Canada, Chile and Brazil. Here country of the first detection is not correlating with the country of maximum prevalence, indicating the spread of the recombinant lineage beyond the country of origin. These lineages could have either emerged in the country where they were first detected but had limited circulation, or country of maximum prevalence where it was not initially detected. Of the other three recombinant lineages detected before the omicron wave, XA was only detected in the UK, while XB emerged in the USA where it was first detected and circulated in maximum prevalence with spread limited to North and Central America (15). In the case of XC, although more than 90% of sequences were detected in Japan, its first sequence was collected from the US. Here even, the country of the first detection is different from the country of maximum prevalence.

### The majority of SARS-CoV-2 recombinant lineages belong to Omicron monophyletic group

To understand the genealogical distribution of recombinant lineages, a phylogenetic analysis was performed [FIG3]. Maximum-likelihood phylogenetic tree of SARS-CoV2 lineages was inferred using lineage representative sequences rooted in the A lineage, which is the closest relative to RatG13 (29). To analyse if any of these 31 recombinant lineages are phylogenetic duplicates assigned with different names, other than analysing one sequence each in the case of non-recombinant lineages, 3 representative sequences each of recombinant lineages were analysed in the phylogenetic tree. Here we could identify, intra-recombinant lineage sequences of all the 31 recombinant lineages clustering together to be genealogically close to each other than any other lineage. We could observe intra-recombinant lineage sequences to be phylogenetically closer to each other than inter-recombinant lineage sequences, validating all the 31 recombinant lineages to be unique lineages resulting from exclusive recombination events.

**Fig 3:**
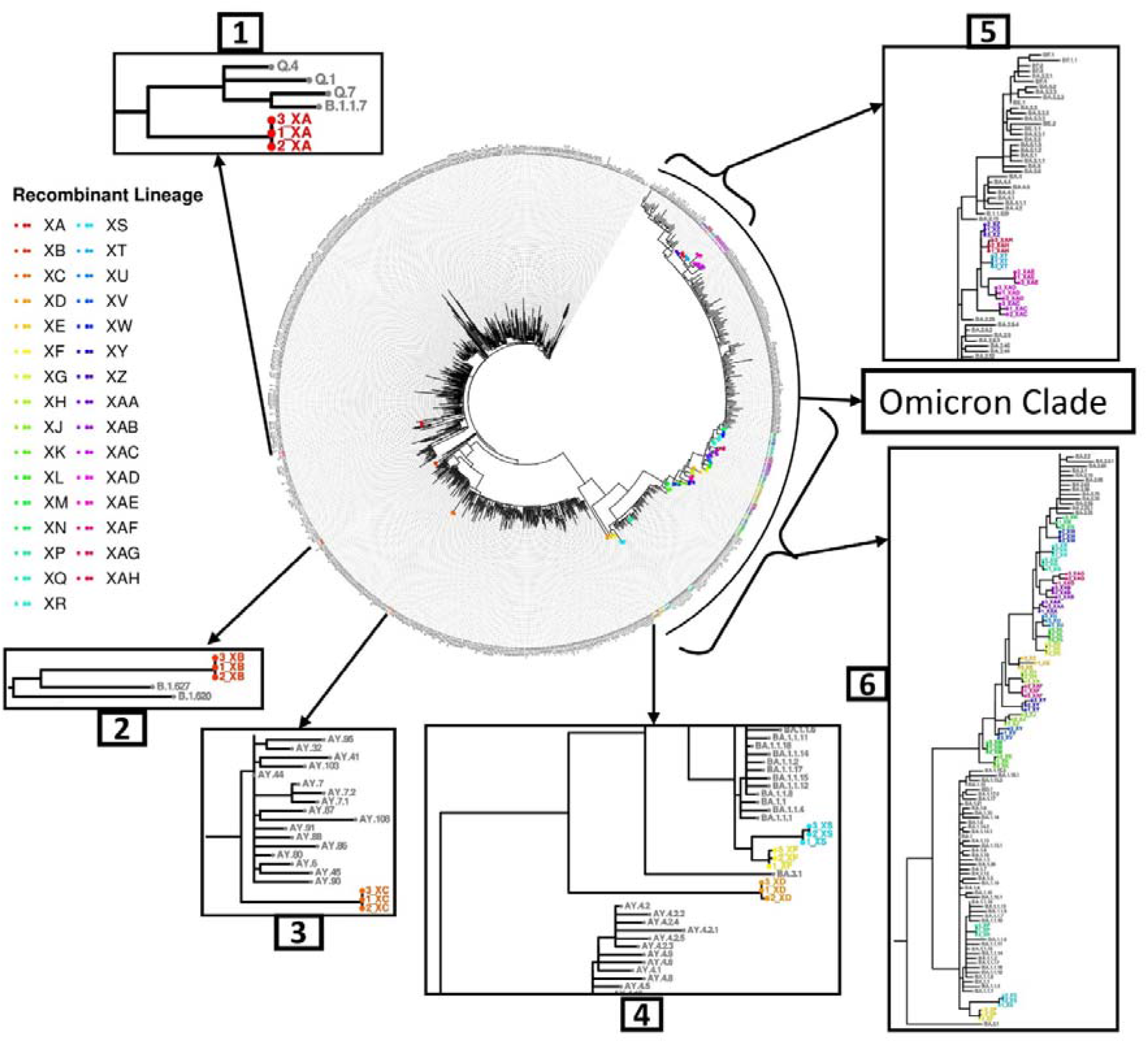
Phylogenetic tree of SARS-CoV2 and genealogical distribution of recombinant lineages: Phylogram in circular layout at the centre represents consensus phylogenetic tree of SARS-CoV2 with three sequences each of recombinant lineages and one sequence each of other lineages. Tip labels are lineage names, with the number of the sequence mentioned followed by an underscore used before the lineage name in the case of recombinant lineage sequences. Recombinant lineage sequence tip labels and tip points are labelled in colours other than black (Legends positioned to the left), while all other lineage sequences are coloured black. 5 insets show zoomed portions of the phylogenetic tree in a rectangular layout. Inset 1 – Shows XA recombinant lineage sequences and all lineage representative sequences sharing the same parent node. Inset 2 – Shows XB recombinant lineage sequences and all lineage representative sequences in two immediate consecutive parent nodes. Inset 3 – Shows XC recombinant lineage sequences with some lineage representative sequences sharing immediate consecutive parent nodes. Inset 4 – Shows XD recombinant lineage sequences with part of lineage representative sequence clade sharing the same parent node. Inset 5 – Shows XT, XZ, XAC, XAD, XAE and XAH, and neighbouring lineage representative sequences. Inset 6 – Shows XF, XS, XP, XK, XM, XV, XJ, XJ, XY, XAF, XH, XE, XG, XL, XU, XAA, XAB, XAG, XQ, XR, XW and XN, and neighbouring lineage representative sequences.

The XA recombinant lineage aligned neighbouring alpha variant clade sharing a common parent node [Fig3: Inset 1]. The XB recombinant lineage near B.1.627 lineage sharing a common node [Fig3: Inset 2]. The XC recombinant lineage aligned near delta variant lineages sharing a common parent node to delta variant subclade [Fig3: Inset 3]. While XD recombinant lineage aligned near omicron lineages sharing a common parent node to omicron monophyletic group [Fig3: Inset 4]. All the remaining 27 of the total 31 recombinant lineages were present in Omicron’s clade. XS, XF and XP were all present inside omicron’s subclade BA. 1 of which, XS and XF were sister groups sharing a common divergence point, with XS being genetically distant [Fig 3: Inset 6]. XZ, XAH, XT, XAE, XAD and XAC together form a subclade in the omicron monophyletic group [Fig 3: Inset 5]. This subclade and BA.2.29 lineage share a common parent node and are in the BA.2 subclade. The rest of the 18 recombinant lineages aligned themselves between BA.1 omicron sub-clade and BA.2 omicron sub-clade [Fig 3: Inset 6]. When arranged in the order of closeness to BA.2, they are XN, XW, XR, XQ, XAG, XAB, XAA, XU, XL, XG, XE, XH, XAF, XY, XJ, XV, XM and XK with former being closer to BA.2 sub-clade and latter being closer to BA.1 sub-clade.

### The majority of SARS-CoV-2 recombinant lineages emerged through recombination between parents of the Omicron variant type

The next step was to understand parent lineages which recombine to form recombinant lineages. Clues regarding the clades in which potential parent lineages could be present were available (**cov-lineage.org**). We identified the most likely parent lineages and the corresponding recombination breakpoint regions in their genomes according to 3SEQ for all the recombinant lineages, except for XB, XP and XW for which the data were insufficient [FIG 4A; S.Table 1]. Both 3SEQ, as well as RDP5 using the default settings, were unable to detect recombination events on these three lineage sequences. Parent sequences of XB were previously reported to be B.1.631 and B.1.634 (15). Of the three recombinant lineages that emerged before the omicron wave, XA is recombinant of the alpha variant sub-lineage (Q4) and B.1.177.18, while XC recombinant of an alpha variant sub-lineage (B.1.1.7) and a delta variant sub-lineage (AY.44). Remaining all 26 recombinant lineages originated from parents of omicron lineages. Among them XD, XF and XS were co-parented by sub-lineages of omicron subclade BA.1 and delta variant sub-lineages. Of the remaining recombinant lineages, 23 were co-parented by BA.1 sub-lineages and BA.2(stealth omicron) sub-lineages, both of which are omicron sub-clades. Summing up, 2 recombinant lineages (XA, XC) were co-parented by alpha variant sub-lineages, and 4 (XC, XD, XF, XS) were co-parented by delta variant sub-lineages, 26 were co-parented by omicron variant sub-lineages of which 23 had omicron sub-lineages as both the parents. To substantiate the evidence of true parent lineages for each of the recombinant lineages, the circulation time span for each of the recombinant lineages and their corresponding parent lineages were analysed [FIG4A]. We observed significant overlap between recombinant lineage time spans and corresponding parent lineage time spans further substantiating the authenticity of the parent lineages. Mosaic structures in each recombinant lineage genome with nucleotide positions inherited from parent 1, parent 2 and breakpoint regions inferred according to 3SEQ were visualized to understand the genetic makeup of each of the recombinant lineages [FIG4B]. We did not find any obvious pattern in recombination with a minimum length of recombinant segment ranging from 2189 nucleotides in XC recombinant lineage to more than 13123 nucleotides in XV recombinant lineage. We further analysed the single nucleotide polymorphism (SNP) patterns with respect to Wuhan Hu 1 strain as a reference, in recombinant and corresponding parent lineages to identify SNPs inherited from each parent [S2A]. We identified the mosaic structure of SNPs in the recombinant lineages correlating with the corresponding identified parent lineages validating recombinant lineages and corresponding parent lineages. To understand if there were recombination hotspots in the genome, where most recombination breakpoint regions fall, and if increased recombination with the advent of omicron variant was due to the development of a recombination hotspot in the genome, recombination breakpoint regions of all the recombinant lineages across the genome were analysed [Fig4C]. Recombination breakpoint 1 was observed to be spread across the genome predominantly in the ORF1ab region, with no specific pattern or hotspots detected. But when we survey recombination breakpoint 2 of all recombinant lineages, even though it is outside the ORF1ab region,60 per cent of it is observed in the 3’UTR region [Fig4C]. Next, we sought to understand if any region of a particular parent lineage or variant was preferably enriched in specific regions of the genome in recombinant lineages [S3B]. Parent variant percentage distribution at each nucleotide position in the genome of recombinant lineages was analysed. We observed more than 75 per cent of recombinant lineages inherited spike sequences from BA.2 sub-lineage parents [S2B]. The overall majority of the recombination events took place among the same variants, with a recombination hotspot in 3’ UTR and potential enrichment of Spike from BA.2 sub-lineage.

**Figure 4:**
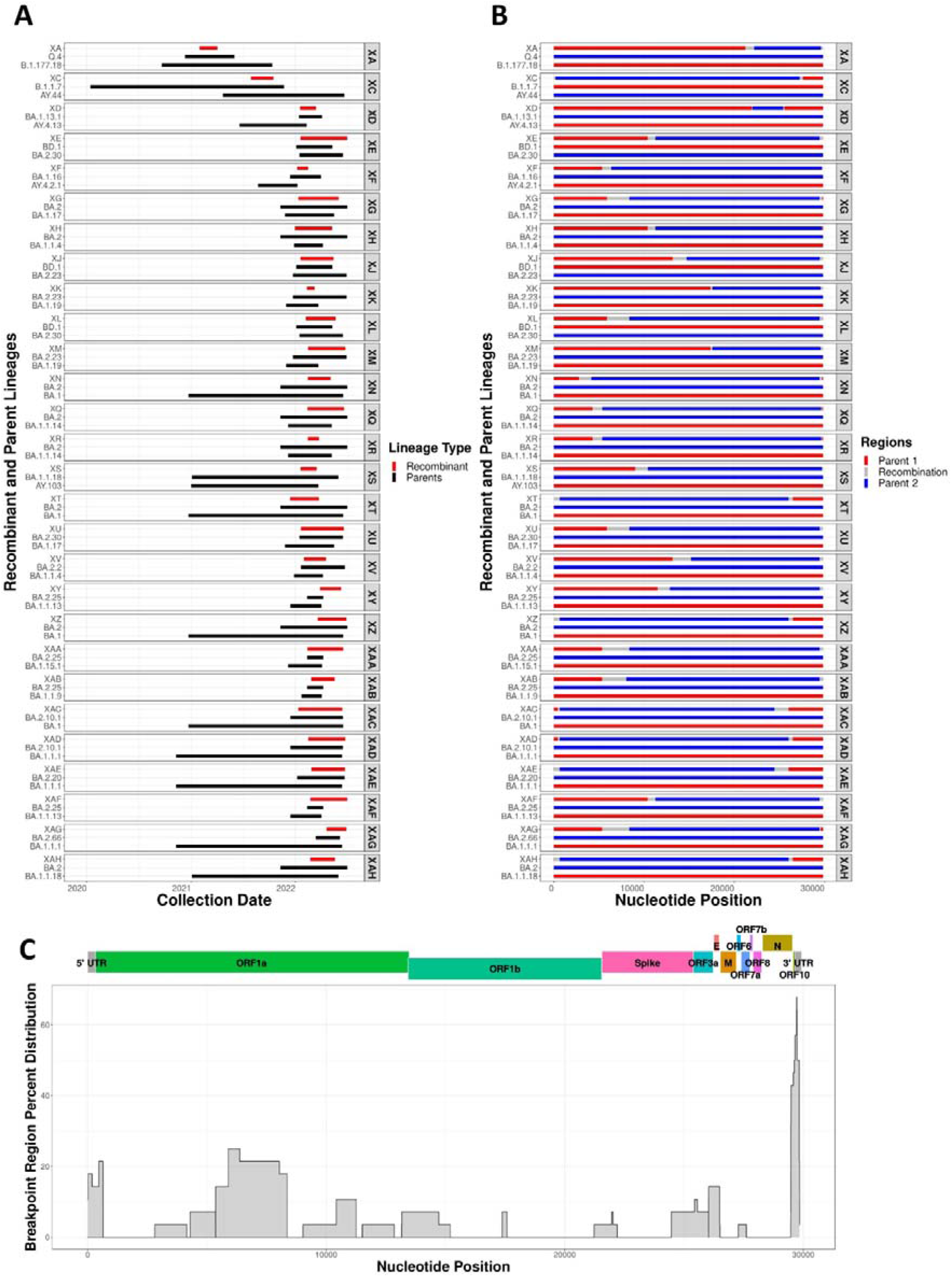
Co-circulation detection, mosaicism and breakpoint region distribution of recombinant lineages with best-identified parent lineages using 3SEQ: (A) Facetted timeline of recombinant lineages and corresponding parent lineages, where each facet representing a recombinant lineage with the facet mentioned to the right side of each facet. The Common X-axis shows the collection date of sequences. Each facet has a unique Y axis having recombinant and parent lineages, with recombinant lineages on top. Horizontal bars indicate the time span between the first and last detection of the corresponding lineage. Red bars indicate recombinant lineages and black bars for parent lineages. (B) Facetted mosaic structure representation of recombinant lineage genomes and corresponding parent lineages with common X-axis showing nucleotide sequence position in SARS-CoV2 genome. Y-axis and facets remain the same as in Fig4A. Coloured segments of red and blue indicate regions from each of the parents, with grey segments representing breakpoint regions predicted by 3SEQ. (C) Percentage of breakpoint regions falling in each nucleotide position of all recombinant lineage genomes with X-axis showing nucleotide position in the genome, Y axis showing the percentage of breakpoints detected in corresponding nucleotide, ORF regions and UTRs are mapped onto the genome and marked on top with dark grey segments indicate 5’ UTR and 3’ UTR regions while different colours showing different ORFs with names marked including ORF1a, ORF1b, Spike, ORF3a, E(Envelope), M(Membrane), ORF6, ORF7a, ORF7b, ORF8, N(Nucleocapsid Protein) and ORF10.

### Comparative analysis of the nucleotide and amino acid sequences of SARS-CoV-2 recombinant lineages

Next, we examined nucleotide sequence variations between genomes of different SARS-CoV2 recombinant lineages [S5A]. We identified differential frequencies of nucleotide polymorphic sites in different regions of the genome ranging from 4.6 sites/100 nucleotide positions in ORF8 to 0.41 sites per 100 nucleotide positions in ORF1b. In the case of spike protein, we observed an intermediate 2.56 polymorphic sites per 100 nucleotide position with 98 inter-recombinant lineage polymorphic sites [S5B]. Compared to that, in the spike protein of recombinant lineages formed post omicron emergence, polymorphic site frequency is reduced to 1.72 polymorphic sites per 100 nucleotide position with 66 nucleotide polymorphic sites. The sense nucleotide variations leading to amino acid changes are the primary driver of viral evolution. To understand the amino acid level changes introduced through recombination events, we analysed inter-recombinant lineage amino acid polymorphism in all the 12 ORFs [Fig5A]. We identified alternative stop codon positions in ORF8 of some recombinant lineages, where early stop codons were identified. Truncated ORF8 proteins having early stop codons are reported in some other lineages of SAR-CoV2 (30). Like nucleotide polymorphic site frequency, we observed a differential number of amino acid polymorphic sites in different ORFs ranging from 7.37 sites per 100 amino acid positions in ORF8 protein to 0.7 sites/100 amino acid positions in ORF1b.

**Figure 5:**
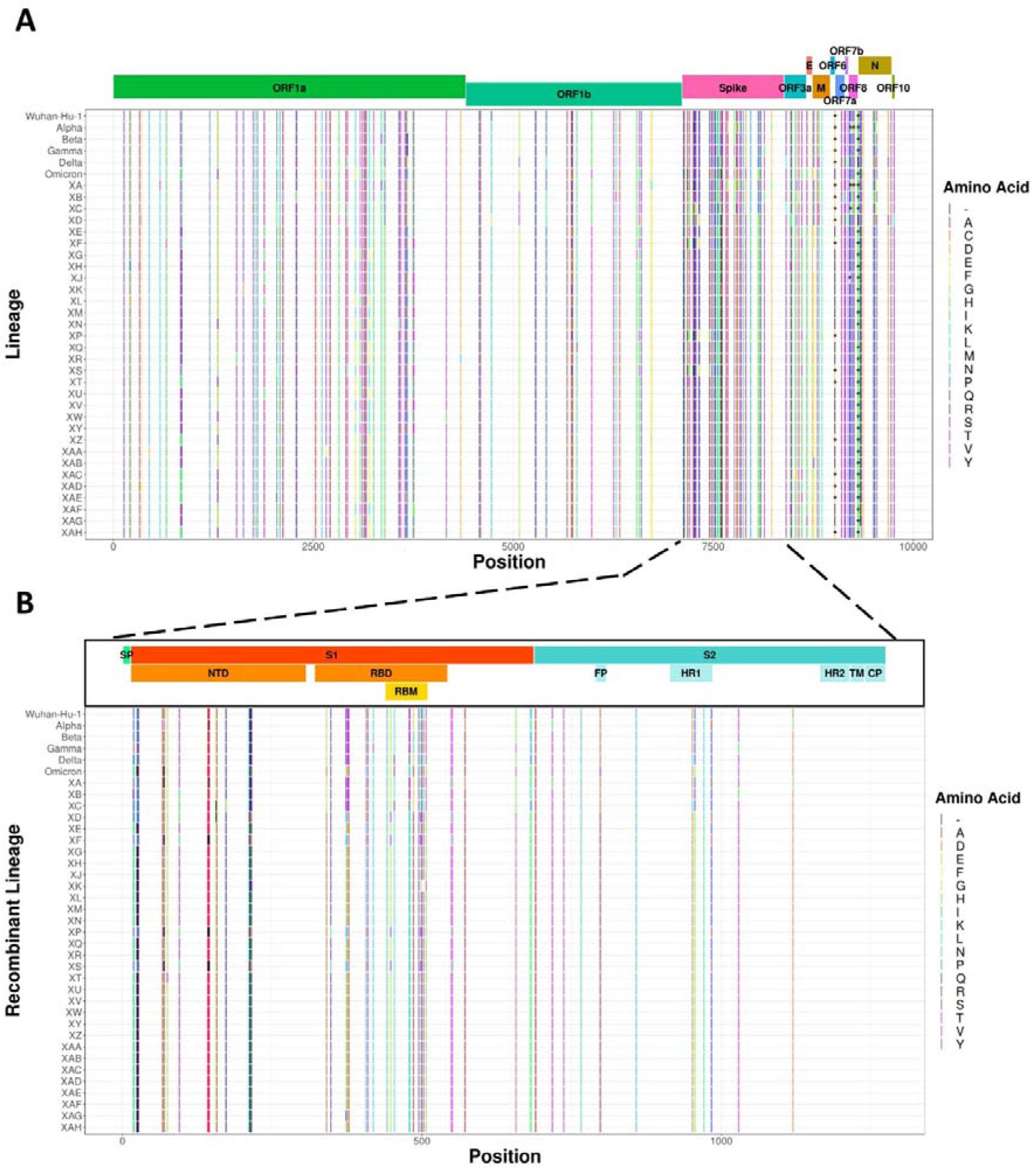
SARS-CoV2 proteome amino acid inter-recombinant lineage polymorphic sites with the spotlight on spike protein: (A) SARS-CoV2 proteome polymorphic amino acid positions are marked using “|”, with black indicating a gap in the amino acid position and different colours representing different amino acids (Colour legends mark amino acids with one letter amino acid codes).”*” represents stop codons. Corresponding amino acids in the inter-recombinant lineage polymorphic sites of Wuhan-Hu-1 strain, Alpha variant(B.1.1.7 lineage), Beta variant(B.1.351 lineage), Gamma variant(P.1 lineage), Delta variant(B.1.617.2 lineage) and omicron variant(B.1.1.529 lineage) are included for comparison. Each of the 12 ORF regions was mapped and marked on top in different colours including ORF1a, ORF1b, Spike, ORF3a, E(Envelope), M(Membrane), ORF6, ORF7a, ORF7b, ORF8, N(Nucleocapsid Protein) and ORF10 (B) SARS-CoV2 Spike protein inter-recombinant lineage amino acid polymorphic sites with both X and Y axis remaining same as Fig5A Different colours representing different amino acids and black indicating a gap in the amino acid position(Colour legends mark amino acids with one letter amino acid codes). Spike ORF sub-regions were mapped and marked on top. Regions marked include SP(Signal Peptide), S1, S2, NTD(N-Terminal Domain), RBD(Ribosome Binding Domain), RBM(Ribosome Binding Motif), FP(Fusion Peptide), HR1 (Heptad Repeat 1), HR2(Heptad Repeat 2), TM(Transmembrane region) and CP(Cytoplasmic region). Different colours mark different spike subregions with S1 and subregions represented in shades of orange, while S2 and sub-regions are represented in shades of cyan.

Spike protein exhibited an amino acid polymorphic site frequency of 1.72 sites per 100 amino acid positions with 64 amino acid polymorphic sites [Fig5B]. Of these 64 sites, 25 of these sites were identified in the N-terminal domain (NTD) region with a frequency of 8.5 polymorphic sites per 100 amino acid positions, 20 of them in the RBD (Receptor Binding Domain) region with 8.9 polymorphic sites/100 amino acid positions, 1 site in fusion protein region with 5.5 polymorphic sites per 100 amino acid positions and 5 sites in Heptad repeat 1 with a frequency of 6.8 polymorphic sites per 100 amino acid positions. When we compare only among recombinant lineages formed post omicron emergence, there are 34 amino acid polymorphic sites, 22 sites are in NTD with the frequency of 7.48 polymorphic sites per 100 amino acid positions, 9 sites in RBD with 4.03 polymorphic sites per 100 amino acid positions and 1 site in Heptad repeat 1 region with a frequency of 1.37 polymorphic sites per 100 amino acid positions. The recombination events in coronaviruses involve the action of exonuclease (NSP14) and other viral polymerase components. It was previously reported that Nsp14 plays an important role in the recombination of coronaviruses (31). To understand if the increased emergence of recombinant lineages parenting omicron variant sublineages, was due to any specific change in the replication and proof-reading machinery in the parent lineages, we analysed the Nsp14 protein sequence in all the recombinant parent lineages and major VOCs [S4A]. We were able to identify I42V mutation conserved across parent lineages belonging to the omicron variant as well as in the Omicron variant parent lineage (B.1.1.529) [S4B]. As RdRp (Nsp12) and Helicase(Nsp13) play important roles in the replication (32), we analysed both the protein sequences but did not find any specific amino acid conservation pattern in parent lineages from omicron lineage, potentially ruling out their role in heightened emergence of recombinant lineages post-emergence of omicron variant [S5A; S5B]. As Nsp10 plays an important role in Nsp14 function acting as a cofactor, we even analysed its amino acid sequence and here also we did not find any specific mutation conservation pattern in omicron variant recombinant parent lineages [S5C].

### Spike protein of SARS-CoV-2 recombinant lineages has an enrichment of amino acid changes that impart immune escape

Spike protein of SARS-CoV-2 is crucial for immune escape and viral fitness and has shown significant sequence variability, potentially due to immune pressure (3). We examined whether Spike protein of recombinant lineages had any specific pattern of amino acid enrichment, especially in the key functional domains. For this, we calculated the amino acid residue conservation score in the spike protein of parent lineages of omicron recombinant lineages relative to non-recombinant omicron lineages (Relative conservation score) [FIG6A]. We identified 28 residues relatively conserved, of which 16 residues reside in the NTD region of the spike, 9 residues in the RBD region and 1 residue in the HR-1 region of the spike. Analysing the spike protein three-dimensional structure showed most of these conserved residues reside exposed on the surface [FIG6B]. Of the 28 relative conserved residues, 11 of them are different from Wuhan Hu 1 strain reference sequence, indicating although non-recombinant omicron lineages underwent variation at the other 17 sites, they were not selected in the recombinant lineages potentially due to their combinatorial or individual deleterious nature in resulting recombinant viruses. Now we sought to understand the significance of all these 11 specific conserved mutated positions in spike protein different from the Wuhan Hu 1 reference strain [Table 1]. 6 of these conserved amino acid positions namely 19, 24, 25, 26, 27 and 213 are located in NTD region while rest 5 of them namely 371, 376, 405, 408 and 493 are located in RBD region. All conserved mutations in the NTD region of I19T, del24-26+A27S and V213G are reported to cause significant evasion from neutralisation antibodies (nAbs) targeting the NTD. Similarly, all conserved mutations in the RBD region namely S371F, T376A, D405N, R408S and Q493R play significant roles in vaccine evasion, broad sarbecovirus neutralizing antibodies escape, poor cross-reactivity among SARS-CoV2 lineages, ACE2 competing antibodies escape and resistance to therapeutic monoclonal antibodies against SARS-CoV2 (7, 33–35). Mutation in Q493 residue lineages play a crucial role in immune and vaccine evasion. Analysing these conserved sites in each of the recombinant lineages shows amino acid variation from the conserved residue in primarily 4 lineages in most of the sites, namely XD, XF, XS and XP [Supplementary Table 2]. Out of these, XD, XF and XS only have a single omicron parent which is BA.1 sublineage, and the other parent, a delta variant sublineage. In the case of XP recombinant lineage, tracking the parent lineages still remains a challenge due to insufficient data. For all the other recombinant lineages parented by both omicron parent lineages, namely BA.1 sublineages and BA.2 sublineages, conserved residue remains unchanged in all positions of these recombinant lineages, except in case XAG at spike amino acid position 371.

**Figure 6:**
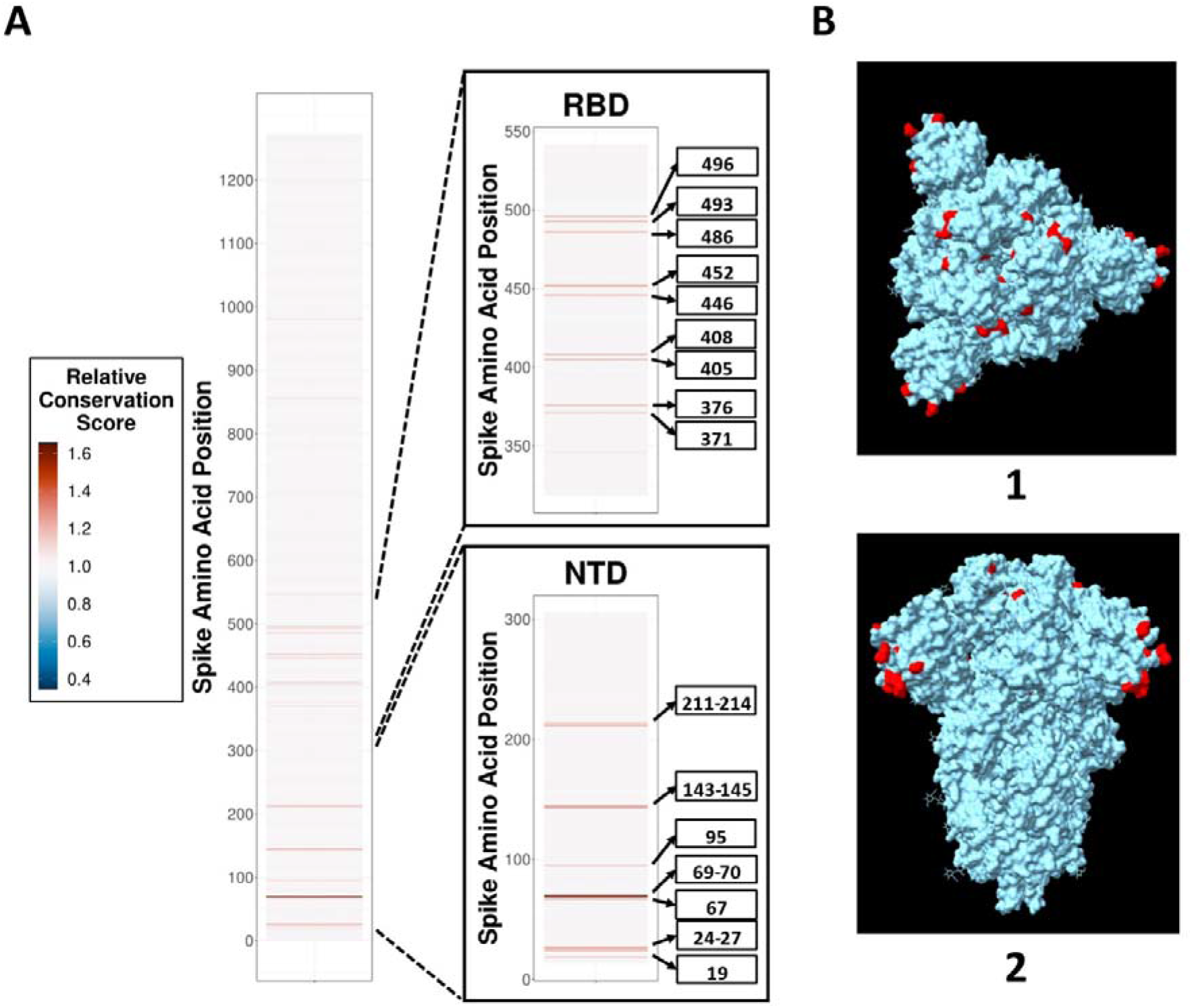
SARS-CoV2 spike relative conserved residues in recombinant lineages and their structural visualization: **(A)** Relative residue conservation score of recombinant lineages with at least one omicron variant parent, relative to non-recombinant omicron variant lineages with red coloured residues relatively more conserved among recombinant lineages than non-recombinant omicron variant lineages. There is only Y-axis here which shows the amino acid position in spike protein. There are two insets: Inset RBD – Relative residue conservation score of RBD zoomed in, with residues having more than 1.1 relative residue conservation score named; Inset NTD – Relative residue conservation score of NTD zoomed in, with residues having more than 1.1 relative residue conservation score marked and named. **(B)** 3D structure of spike glycoprotein trimer in prefusion closed configuration(PDB ID–6VXX) with red coloured residues showing conserved amino acid positions in recombinant lineages with at least one omicron variant parent relative to non-recombinant omicron variant lineages. There are 2 insets: Inset 1 – Top view of the spike; Inset 2 – Side view of the spike.

**Table 1:** SARS-CoV2 spike relative conserved mutated positions in recombinant lineages with discovered relevance in viral transmission and immune escape. Column 1: SARS-CoV2 spike relative conserved mutated positions in recombinant lineages; Column 2: Wuhan Hu 1 strain reference sequence residue at the conserved positions; Column 3: Amino acid residue conserved among recombinant lineages(with at least one omicron variant parent lineage) at that relative conserved positions; Column 4: Relative conservation score of each conserved position; Column 5: Reported significance of the conserved mutation in the relative conserved position; Column 6: References reporting the significance of the conserved mutations in the relative conserved positions.

| Amino acid Position | Wuhan Reference Residue | Recombinant Conserved Residue | Relative Conservation Score | Significance | Reference |
| --- | --- | --- | --- | --- | --- |
| 19 | T | I | 1.18 | T19I: Significant evasion from NTD-targeted neutralizing antibodies (nAbs) | (34) |
| 24 | L | - | 1.18 | del24-26+A27S: Loss in neutralization activity of NTD-directed monoclonal antibodies(mAbs) | (34) |
| 25 | P | - | 1.18 | del25-27 : Significant evasion from NTD-targeted neutralizing antibodies ;<br>del24-26+A27S: Loss in neutralization activity of NTD-directed monoclonal antibodies | (33, 34) |
| 26 | P | - | 1.18 | del25-27 : Significant evasion from NTD-targeted neutralizing antibodies ;<br>del24-26+A27S: Loss in neutralization activity of NTD-directed monoclonal antibodies | (33, 34) |
| 27 | A | S | 1.23 | A27S: Reduce spike sensitivity to neutralization by sera from BNT/BNT vaccinated individuals;<br>del24-26+A27S: Loss in neutralization activity of NTD-directed monoclonal antibodies | (34, 35) |
| 213 | V | G | 1.13 | V213G: Reduce spike sensitivity to neutralization by sera from BNT/BNT vaccinated individuals | (35) |
| 371 | S | F | 1.13 | S317F: Induce large-scale escapes of broad sarbecovirus neutralizing antibodies ;<br>Reduce spike sensitivity to neutralization by BNT/BNT sera in the range of 2 to 5 fold | (7, 35) |
| 376 | T | A | 1.18 | T376 mutation helps ACE2 competing antibodies escape | (7) |
| 405 | D | N | 1.17 | D405N: Significant escape of BA.1 lineage omicron-specific neutralizing antibodies ;<br>Induce large-scale escapes of broad sarbecovirus neutralizing antibodies ;<br>D405 mutation helps ACE2 competing antibodies escape ;<br>Alters the antigenic surface that disrupts the binding of antibodies;<br>The main reason for poor crossreactivity among BA.2/BA.3/BA.4/BA.5 sublineage. | (7) |
| 408 | R | S | 1.2 | R408S: Induce large-scale escapes of broad sarbecovirus neutralizing antibodies<br>R408 mutation helps ACE2 competing antibodies escape ;<br>Alters the antigenic surface that disrupts the binding of antibodies; | (7) |
| 493 | Q | R | 1.23 | Q493R: Emerges during bamlanivimab/etesivimab cocktail treatment ;<br>Causes resistance to bamlanivimab and etesivimab ;<br>Q493 is critical for binding to Class 2 and 3 antibodies ;<br>Q493 mutations increase binding affinity to the ACE2 | (36) |

## DISCUSSION

Genetic recombination is known to occur at different rates among RNA virus families and plays a crucial role in viral evolution, emergence and epidemiology (19). It has been reported to contribute to altered viral host-tropism, enhanced virulence, host immune evasion and development of resistance to antivirals (37). It is very common for Retroviruses and other positive sense RNA viruses and is rarely observed in the case of negative sense RNA viruses. In the case of HIV, genetic recombination has contributed to the emergence of highly prevalent recombinant forms with improved viral fitness(18). Similarly, in the case of HCV, recombinant lineages have been reported to circulate widely for a prolonged period of time (38). Among Sarbecoviruses, recombination is commonly observed, although at varying rates (24, 29). For seasonal human coronaviruses (HCoVs) 229E, HKU1, NL63 and OC43, frequent recombination is observed between individual genomes and rarely between different clades (24). It is considered a key contributor to the emergence of new HCoVs including SARS-CoV, MERS and SARS-CoV-2 (22). Genetic recombination in RNA viruses is akin to sexual reproduction where a chimeric progeny is generated with shared genetic features from parental strains. At the molecular level recombination requires genetic sequence similarity between parents, which allows template switching by the viral RNA polymerase during viral genome replication. This requires co-infection of the host cell with both parental strains, which are usually different lineages of the same virus or related viruses, are co-circulating in the same location and present within the same host (37). In our analysis, we have demonstrated that increased recombination events observed during the Omicron wave were not due to higher genomic sampling or co-circulating genetic diversity (Supplementary Fig 2B). We observed that the majority of recombinant lineages emerged in Northern America and Europe. These geographical regions comprise global travel hotspots, which could contribute to the introduction and co-circulation of multiple SARS-CoV-2 lineages (39), which in turn could undergo recombination. Prolonged infection of immunocompromised individuals can lead to the emergence of new SARS-CoV-2 variants and has been postulated as responsible for the emergence of Omicron VOC in south Africa (40). Similar events could also underlie the emergence of recombinant lineages from the individuals with prolonged co-infection with parent strains. RNA viruses replicate at a rapid rate and have a large population size to maintain genetic diversity which is necessary to overcome selection pressure. As a mechanistic by-product of rapid and error-prone replication, they can accumulate deleterious mutations. Recombination is an evolutionarily conserved mechanism through which RNA viruses purge deleterious mutations (37). Acquisition of advantageous genetic features through recombination is rarely observed in RNA viruses (37). Interestingly in our study, we observed genetic fixation of amino acid residues in the key domain of Spike protein of recombinant Omicron lineages, which can facilitate immune escape.

The spike protein of SARS-CoV-2 is the key determinant of viral tropism, transmission and pathogenesis (41, 42). It is also the primary target of host immune response, especially antibody-mediated neutralization (43). Spike protein binds to the ACE2 receptor on the host cell surface through its receptor binding domain, which is the main target of neutralizing antibodies (43). Other domains of Spike, such as the N terminal domain and heptad repeats are also functionally important and targeted by the host antibodies (44, 45). Through the course of the COVID-19 pandemic, SARS-CoV-2 has continuously acquired a range of amino acid changes, which have facilitated resistance to or escape from host antibodies (6–12). These mutations accumulated more rapidly in the Omicron VOC, allowing escape from the host and vaccine-mediated immunity and causing widespread infections (6–12). Another altered feature of the omicron Spike has been amino acid changes that strengthened binding to ACE2 receptor and enhanced viral transmission (46). In our analysis, we compared the amino acid conservation in the Spike protein of recombinant Omicron lineage when compared to non-recombinant Omicron lineages. We found a number of amino acids relatively conserved in the recombinant Omicron lineages, especially in the RBD and NTD regions, which have been reported to facilitate escape from neutralizing antibodies (Fig6; Table 1) (7, 34, 35). Some conserved amino acids were also reported to improve ACE2 binding and/or enhance viral transmission (Table 1) (36). These data suggest an active role of accelerated recombination during the Omicron wave, in the selection of amino acids that facilitated escape from host immunity and improved viral fitness. The Spike protein also happens to be a key target of host T-cell mediated immunity (47). It will be interesting to examine whether recombination events contributed to the escape of SARS-CoV-2 from T-cell-mediated cellular immunity as well. A limitation of studies inferring viral evolution based on genomic surveillance data is the bias introduced due to differences in sampling intensity across geographical regions. This applies to SARS-CoV-2 as well, where the disparity in genomic surveillance is obvious (48); hence the conclusions regarding the origin, prevalence and relative frequency of different lineages must be drawn with caution. At the same time, this also highlights the paramount importance of the active genomic surveillance of SARS-CoV-2, in humans as well as animal reservoirs, to understand the drivers and direction of viral evolution.

### Ideas and Speculation

Recombination in RNA virus genomes is executed by viral RNA polymerase, which in turn requires enzymatic assistance of other non-structural proteins. For SARS-CoV-2 the NSP14 exonuclease has been reported to play a key role in genetic recombination (31). In the case of SARS-CoV-2, we observed an amino acid change I42V, which was conserved in Omicron lineage and not found in other VOCs (Sup Figure 4). 3D modelling of the enzyme shows residue 42 is not proximal to the RNA binding pocket. Nevertheless, it will be interesting to examine whether this I42V change has any forbearance on enhanced recombination rates observed among SARS-CoV-2 of Omicron lineages. Another interesting observation of our study is the concentration of recombination breakpoints in the 3’ UTR of the parent lineages. This region serves as the initiation point for viral genome replication and has important regulatory roles in the same (49). It will be important to explore if there are specific sequence and RNA secondary changes in the 3’ UTR of the Omicron lineages SARS-CoV-2, which could be responsible for accelerated recombination. Furthermore, recombination can lead to a genetic shift allowing the virus to jump the host-species barrier between animal reservoirs and humans, resulting in an outbreak (22, 50). It is well established that SARS-CoV-2 has now spread to a range of non-human species, however genomic surveillance in these species is very limited (50, 51). Although the prevalence of recombinant SARS-CoV-2 lineages remains low currently, considering the enhanced frequency of recombination events and expanded host range of SARS-CoV-2, it is a matter of concern vis-à-vis the emergence of new VOCs in future, especially from Zoonotic origin.

## MATERIALS AND METHODS

### Sequences, metadata and protein structure retrieval

#### Recombinant Sequences

All Recombinant lineage sequences with a collection date between 1^st^ November 2019 and 14^th^ July 2022 were retrieved from the GISAID database on 14^th^ July 2022 (26). These were filtered to discard incomplete sequences with lengths less than 29000 nucleotides, gapped sequences having more than 5% ambiguous nucleotide positions, and sequences with no sequence collection date information available. Corresponding sequence metadata information was also retrieved from the same database.

#### Control Sequences

All 1,206,055 complete SARS-CoV2 sequences (Taxid: 2697049) deposited in the NCBI Virus database (27), isolated between 31^st^ October 2019 and 14^th^ July 2022 were retrieved on 14^th^ July 2022 with appropriate filters including 1) genome size should be between 29000 and 31000 nucleotides, 2) sequences with more than 1% of ambiguous nucleotides positions are avoided, 3) sequences isolated from lab hosts are avoided. Corresponding sequence metadata information including geographic location, country of isolation, length of the sequence, collection date of the sequence and pangolin lineage of the sequence were retrieved from the same database.

#### Reference Sequence

Wuhan Hu-1 reference strain genome sequence, all ORF nucleotide sequences and Nsp10, Nsp12, Nsp13 and Nsp14 protein sequences were retrieved from the NCBI database (52).

#### Protein Structure

3-D structures of SARS-CoV2 spike in prefusion closed (PDB ID: 6VXX) and Nsp14 protein in complex with Nsp10 and RNA (PDB ID: 7N0B) were retrieved in .pdb format from Protein Data Bank (PDB) (53–55).

### Analysis, segregation and scoring of sequences

Sequences collected from both GISAID as well as NCBI Virus databases were independently segregated and clustered into separate lineages using the Phylogenetic Assignment of Named Global Outbreak LINeages (PANGOLIN) assignment tool (26, 27, 56, 57).PANGOLIN tool updated to the latest version of v4.1.2 was utilised, with Constellations version v0.1.10 and Scorpio version v0.3.17. PANGOLIN data was updated to the latest release version v1.8. After lineage segregation and clustering, sequences categorised as lineage unassigned by the PANGOLIN tool were discarded.

NextClade with the latest SARS-CoV2 dataset was utilised to score lineage segregated sequences based on overall quality control scores (QC score), both for recombinant as well as a control set of sequences(58) **(https://clades.nextstrain.org**). Overall QC score is an aggregated score compiling individual missing data score, mixed sites score, private mutations score, mutation clusters score, scoring based on premature stop codons and frameshifts score. Sequences with bad and mediocre scores were discarded, and further best three sequences with ‘good’ QC categorisation from each lineage were extracted into separate files. These sequences were further utilised for phylogenetic analysis, spike inter-recombinant mutation mapping and recombination analysis.

### Phylogenetic analysis of the sequences

Wuhan Hu-1 reference strain genome sequence, the best overall QC scoring sequence from each control lineage with recombinant lineages removed and the top 3 overall QC scoring sequences from each recombinant lineage together underwent masking and multiple sequence alignment using the alignment option of the PANGOLIN tool (57). The maximum likelihood tree was inferred using IQ-TREE 2 utilising the GTR+Γ model of nucleotide substitution, minimum branch length of 0.0000000001 nucleotide substitutions persite and ultrafast bootstrapping with 1000 replicates (59, 60). The phylogenetic tree was rooted in the A lineage which is closest to RatG13 (29).

### Identification and Visualisation of Recombination Breakpoints and parent lineages

The top 3 overall QC score sequences from all potential parent lineages and corresponding recombinant lineages together underwent multiple sequence alignment using MAFFT(Multiple Alignment using Fast Fourier Transform) automatic configuration(61). Mosaic structure and recombination breakpoint regions between these aligned sequence’s parent and recombinant lineages were detected using 3SEQ (62). If 3SEQ was unable to find the parent then, potential parent lineages used to find the breakpoint region were sequentially subsampled and breakpoints with those subsampled parents were detected. In cases where 3SEQ failed even after subsampling, then we tried predicting parent and breakpoint regions with RDP5 in the default configuration (63).

Recombination events were visually verified using a snipit (**https://github.com/aineniamh/snipit**). For each recombinant lineage, the top three overall QC scoring sequences from both identified parent lineages, reference (Wuhan Hu-1 strain) genome sequence and top three best QC scoring corresponding recombinant lineage sequences underwent multiple sequence alignment using MAFFT followed by visualisation of single nucleotide polymorphisms relative to the reference sequence using snipit tool(61), (**https://github.com/aineniamh/snipit**). Region inherited from one of the parents in both recombinant lineage sequences as well as that parent sequences were manually marked for each recombinant lineage visualising the recombination event.

### Lineage consensus sequence generation and mapping of inter-lineage sequence polymorphisms

The top 5 overall QC scoring sequences from all lineages (both recombinant and non-recombinant) were extracted and stored in separate files. These sequences were aligned separately using MAFFT in automatic configuration (61). These aligned sequences were trimmed with parameter -gt 0.5 to remove all amino acid positions which are gaps in more than 50 percent of sequences(64). This trimmed sequences underwent consensus sequence generation with gap filling using an in-house generated program scripted in R. Blastn was used to identify each of the ORF sequence start and end points in recombinant lineage consensus sequences(65). These sequences together with Wuhan Hu 1 reference sequence were aligned using MAFFT in automatic configuration and inter-lineage nucleotide polymorphism was identified using Rstudio(61, 66). Scripts for the same are available.

For each of the 31 recombinant lineages and other non-recombinant lineages, the lineage consensus sequence was used as subject sequence libraries, with the Wuhan Hu-1 reference sequence of each ORFs as the query sequence and the following blastn parameters of - subject_besthit, -max_hsp 1, -gap open 0 and -gap extend 0. In each of these blastn output files, one nucleotide position is subtracted from the start position to make it into a bed file suitable for bedtools (67). ORF sequences from each of the 31 recombinant lineages consensus sequences were extracted using bedtools providing the above-generated bed file coordinates. Frameshift was introduced in ORF1b of each of the recombinant lineages.

ORF sequences from each of the lineages and the reference Wuhan Hu 1 ORF sequences were translated into a protein sequence using EMBOSS Transeq tool, multiple sequences aligned using MAFFT in automatic configuration, and inter-lineage amino acid polymorphism identified using Rstudio(61, 66).

### Residue conservation analysis and 3D structural visualisation

Spike sequences from each of the lineages’ consensus sequences were extracted using bedtools providing above generated bed file coordinates. Consensus spike sequence from each of the lineages was translated into a protein sequence using EMBOSS Transeq tool (66). These control lineages spike sequences and recombinant lineages spike sequences were together aligned using MAFFT using automatic configuration and trimmed using trial with parameters -gt 0.5(61). Omicron lineage spike sequences without recombinant lineages and recombinant lineage spike sequences were extracted and split into two respective files. Conservation scores of each residue of spike recombinant lineages with at least one omicron parent relative to omicron were calculated in Rstudio utilising a customised version of conserv function in Bio3D package (68).

3D structure of the spike in prefusion closed structure(PDB ID: 6VXX) and Nsp14 in complex with Nsp10 and RNA(PDB ID: 7N0B) retrieved from PDB were visualised using CHIMERAX (53–55, 69).

### Graphical Visualisation and analysis

All graphical images were plotted and visualised using RStudio. Packages including ggmap (**https://github.com/dkahle/ggmap)**, map (**https://CRAN.R-project.org/package=maps)**, RColorBrewer (**https://CRAN.R-project.org/package=RColorBrewer)**, scales (**https://CRAN.R-project.org/package=scales)**, gridExtra (**https://CRAN.R-project.org/package=gridExtra)**, scatterpie (**https://CRAN.R-project.org/package=scatterpie)**, ggtree, treeio, tidyverse, treedataverse, writexl (**https://CRAN.R-project.org/package=writexl)**, seqinr (**https://CRAN.R-project.org/package=seqinr)**, zoo (**https://CRAN.R-project.org/package=zoo)** and Biostrings (**https://bioconductor.org/packages/Biostrings)** were utilised for data analysis and visualisation.Scripts for the same are available **(70–72)**

## Acknowledgments

We gratefully acknowledge all data contributors, i.e., the Authors and their Originating laboratories responsible for obtaining the specimens, and their Submitting laboratories for generating the genetic sequence and metadata and sharing via the GISAID Initiative, on which this research is based. 3D structural visualisations performed using UCSF ChimeraX, developed by the Resource for Biocomputing, Visualization, and Informatics at the University of California, San Francisco, with support from National Institutes of Health R01-GM129325 and the Office of Cyber Infrastructure and Computational Biology, National Institute of Allergy and Infectious Diseases. This work is supported by research grants from Crypto Fund, SERB, and DBT-BIRAC to ST Lab. RS is supported by fellowship support from CSIR.

## SUPPLEMENTARY FIGURES & TABLES

**Supplementary Figure 1:**
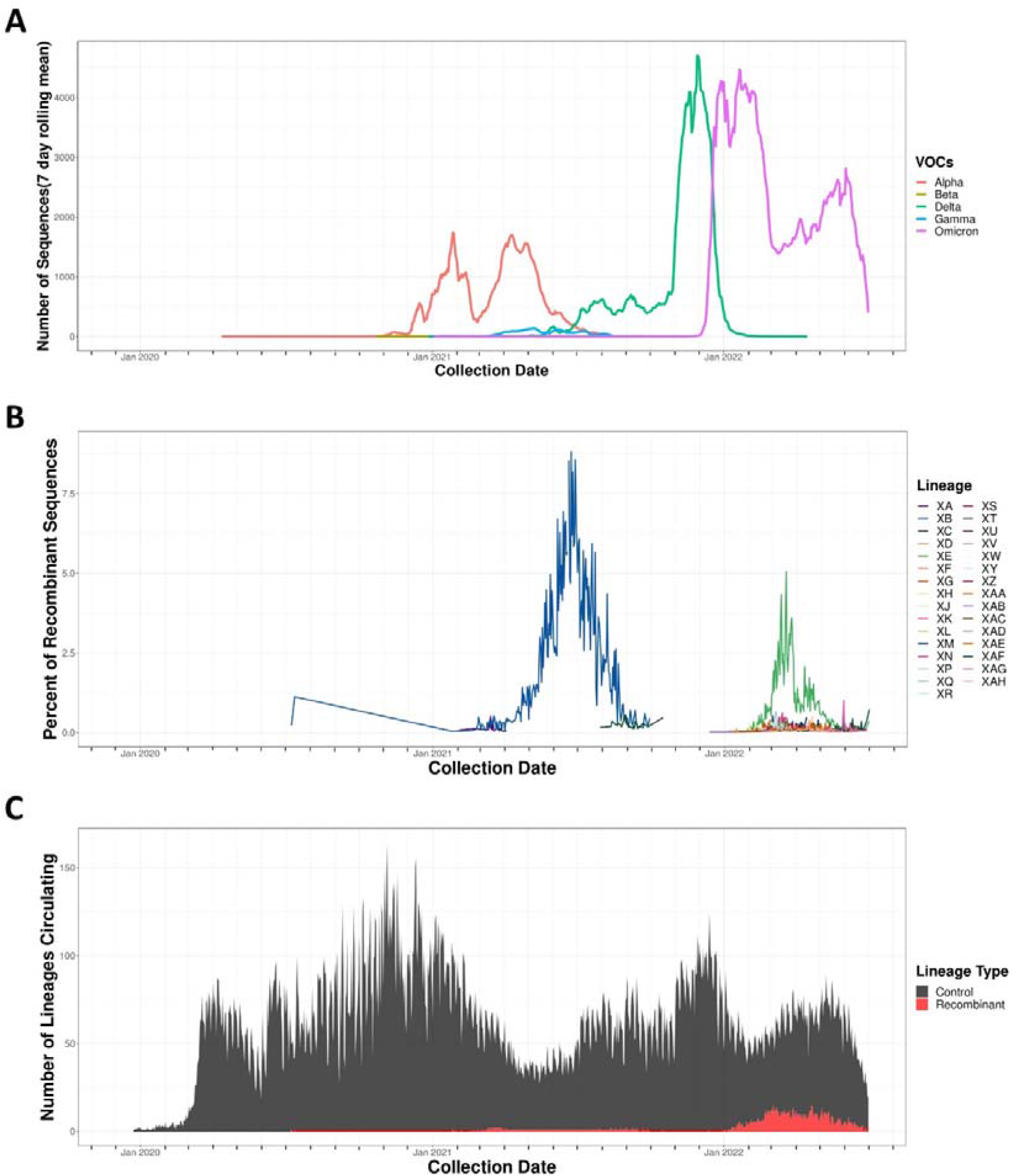
Temporal Distribution of SARS-CoV2 VOCs sequences, percentage temporal distribution of SARS-CoV2 recombinant lineage sequences and a daily number of lineages circulating: (A) Prevalence of SARS-CoV2 variants of concerns(VOC) with X-axis showing date of sequence collection, Y-axis showing several sequences collected and deposited in 7 days rolling average. Different VOCs are represented in unique colours. (B) Percentage of sequences belonging to each recombinant lineage with X-axis and Y-axis same as that of Supplementary Figure 1A. Different recombinant lineages are represented in unique colours which is the same as that used to represent recombinant lineages in FIG1A. (C) Area curve showing the number of unique lineages prevalent daily, with the X-axis representing the collection date of the lineage sequence, the Y-axis representing the number of unique lineages circulating each day and colours representing the type of lineage(recombinant – red, all lineages – dark grey. X axis of all three graphs ranges from 1^st^ December 2019 to 1^st^ July 2022 with axis ticks indicating months and are aligned making them temporally comparable to each other.

**Supplementary Figure 2:**
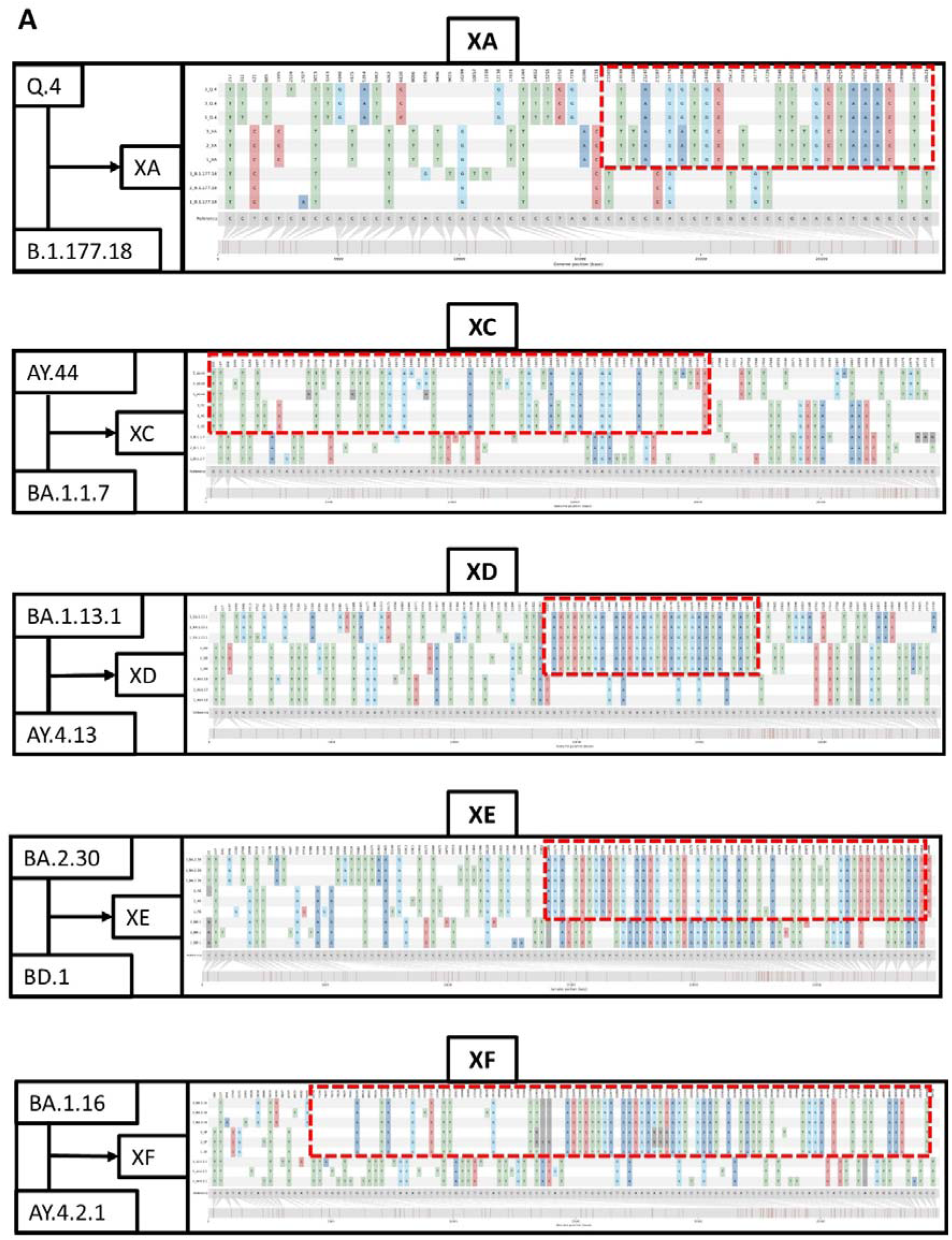
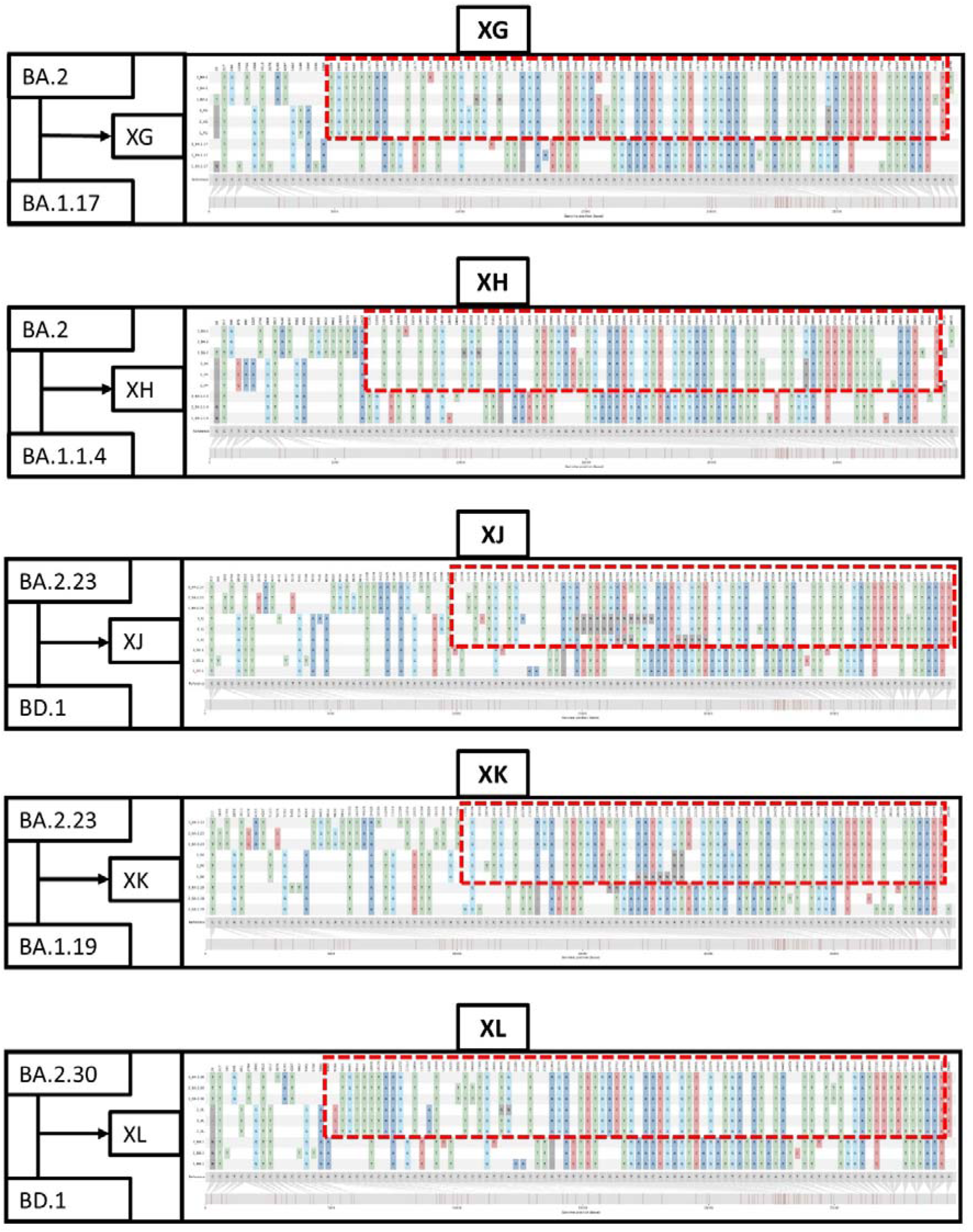
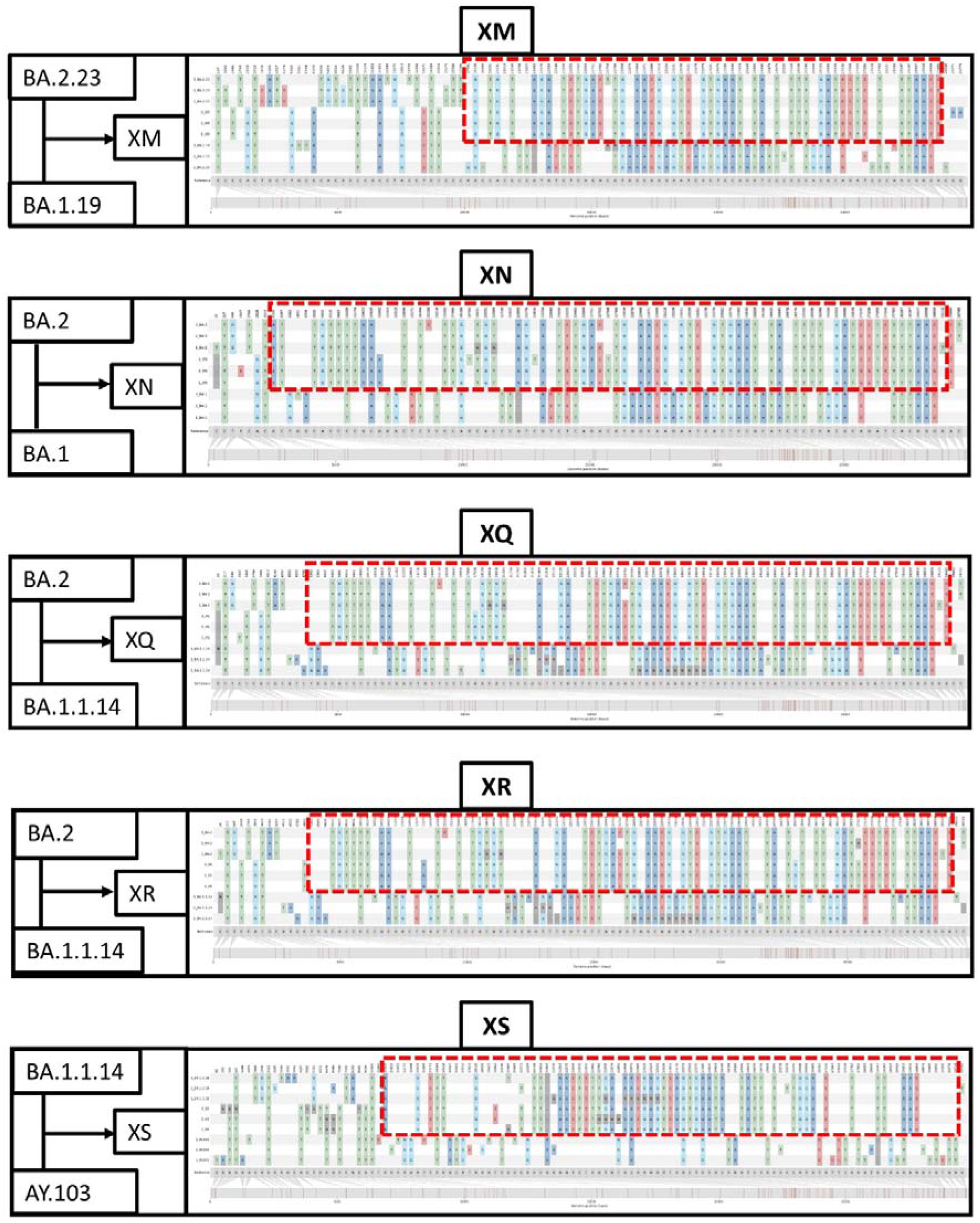
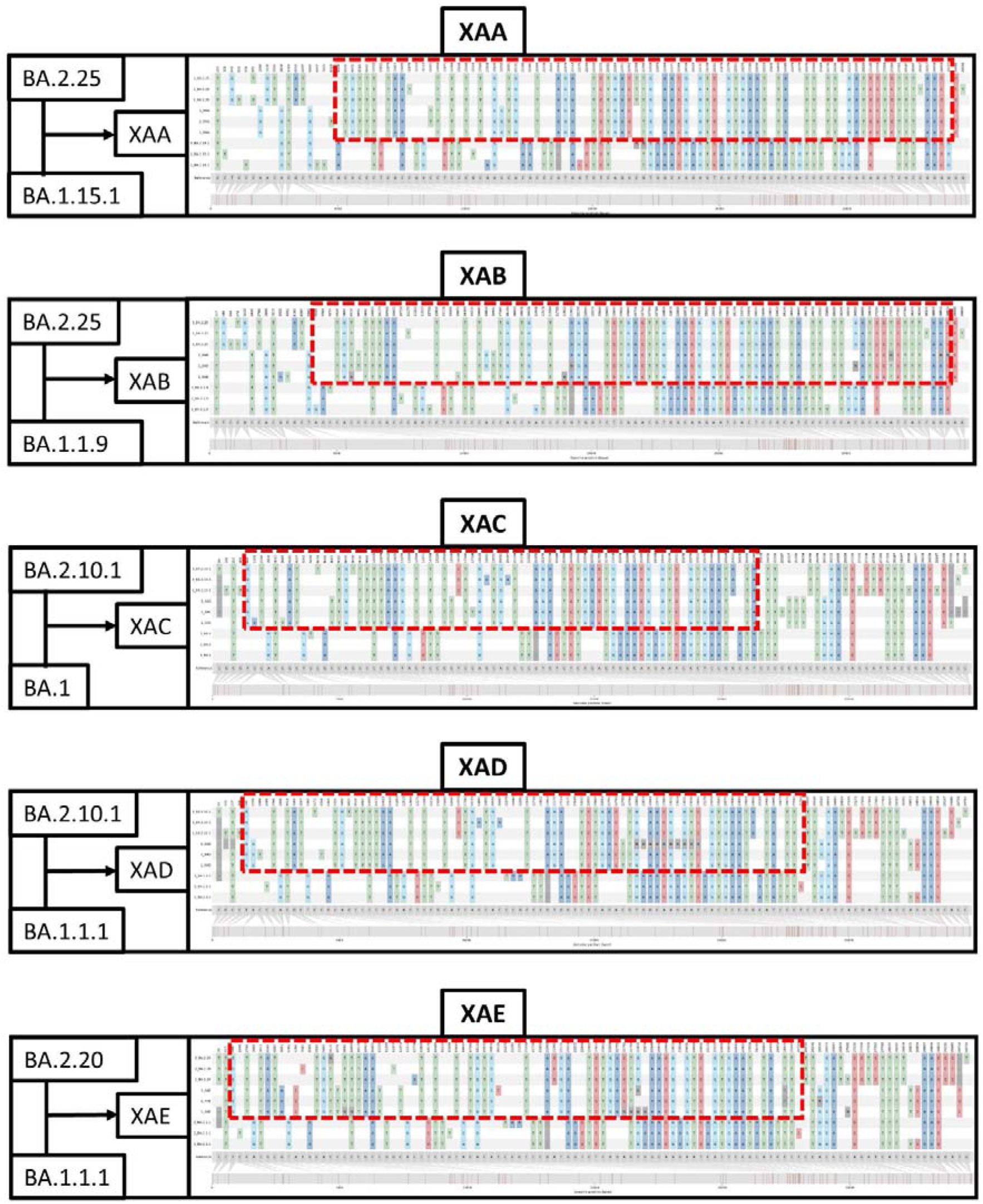
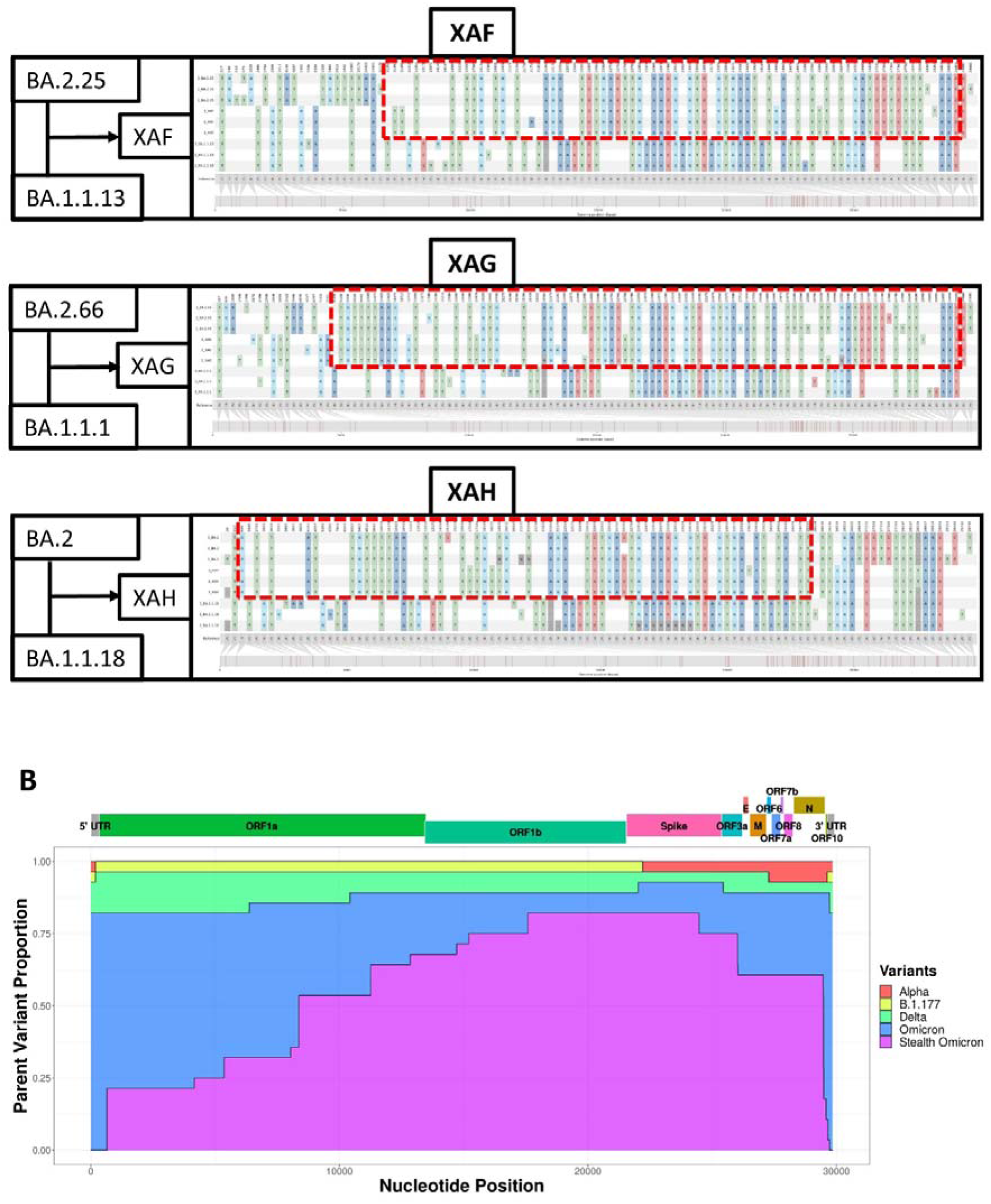
Genome Single Nucleotide Polymorphisms distribution of recombinant lineages and their identified parental lineages showing mosaicism and parent variant proportion distribution in recombinant lineages: (A) Single nucleotide polymorphisms patterns in recombinant lineage sequences and parental lineage sequence with respect to reference Wuhan Hu 1 strain nucleotide sequence using snipit. Top three sequences represent parent 1, middle three sequences belong to corresponding recombinant lineage sequence, and bottom three sequence are parent 2 lineage sequences. Red rectangular box in each image with broken lines represents minimum region in recombinant lineage inherited from parent 1 as predicted by 3SEQ with rest of the recombinant lineage region inherited from parent 2 lineage. Black box to the right have names of the parent and recombinant lineages mentioned in text boxes with arrows directing from parents to recombinant lineage. (B) For each nucleotide position in the genome, proportions of recombinant lineages inheriting that nucleotide position from each parent lineage are represented in a stacked area curve. X axis shows genome nucleotide position, Y axis shows parent variant proportion distribution. Different colours represent the proportion contributed by different variants While 5’ and 3’ Untranslated Regions (UTRs) are mapped and marked on top in dark grey, each of the 12 ORF regions are represented in different colours including ORF1a, ORF1b, Spike, ORF3a, E(Envelope), M(Membrane), ORF6, ORF7a, ORF7b, ORF8, N(Nucleocapsid Protein) and ORF10.

**Supplementary Figure 3:**
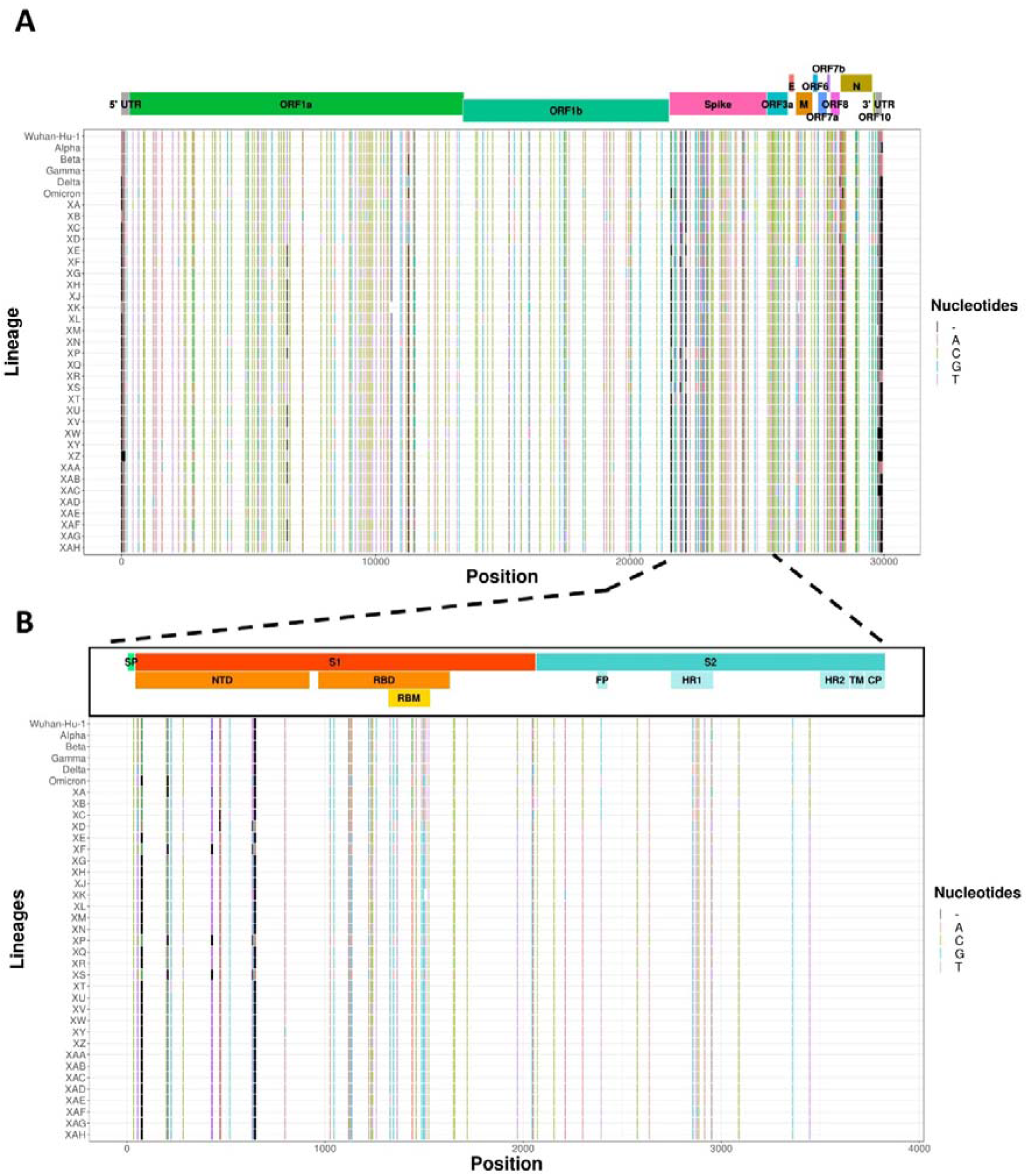
SARS-CoV2 genome inter-recombinant lineage nucleotide polymorphic sites with the spotlight on spike: (A) SARS-CoV2 polymorphic nucleotide positions mapping with X-axis representing nucleotide position in the genome, Y-axis representing lineages, nucleotides in the inter-recombinant lineage polymorphic positions marked using “|”, with black indicating a gap in the nucleotide position and different colours representing different nucleotides (Colour legends mark nucleotides with one letter codes). Corresponding nucleotides in the inter-recombinant lineage polymorphic sites of Wuhan-Hu-1 strain, Alpha variant(B. 1.1.7 lineage), Beta variant(B.1.351 lineage), Gamma variant(P.1 lineage), Delta variant(B. 1.617.2 lineage) and omicron variant(B. 1.1.529 lineage) are included for comparison. While 5’ and 3’ Untranslated Regions(UTRs) are mapped and marked on top in dark grey, each of the 12 ORF regions is represented in different colours including ORF1a, ORF1b, Spike, ORF3a, E(Envelope), M(Membrane), ORF6, ORF7a, ORF7b, ORF8, N(Nucleocapsid Protein) and ORF10. (B) SARS-CoV2 Spike ORF inter-recombinant lineage nucleotide polymorphic sites with both X-axis, Y-axis and colour schemes for nucleotides remaining the same as Supplementary Figure 3A. Spike ORF sub-regions were mapped and marked on top. Regions marked include SP(Signal Peptide), S1, S2, NTD(N-Terminal Domain), RBD(Ribosome Binding Domain), RBM(Ribosome Binding Motif), FP(Fusion Peptide), HR1(Heptad Repeat 1), HR2(Heptad Repeat 2), TM(Transmembrane region) and CP(Cytoplasmic region). Different colours mark different spike subregions with S1 and subregions represented in shades of orange, while S2 and subregions are represented in shades of cyan.

**Supplementary Figure 4:**
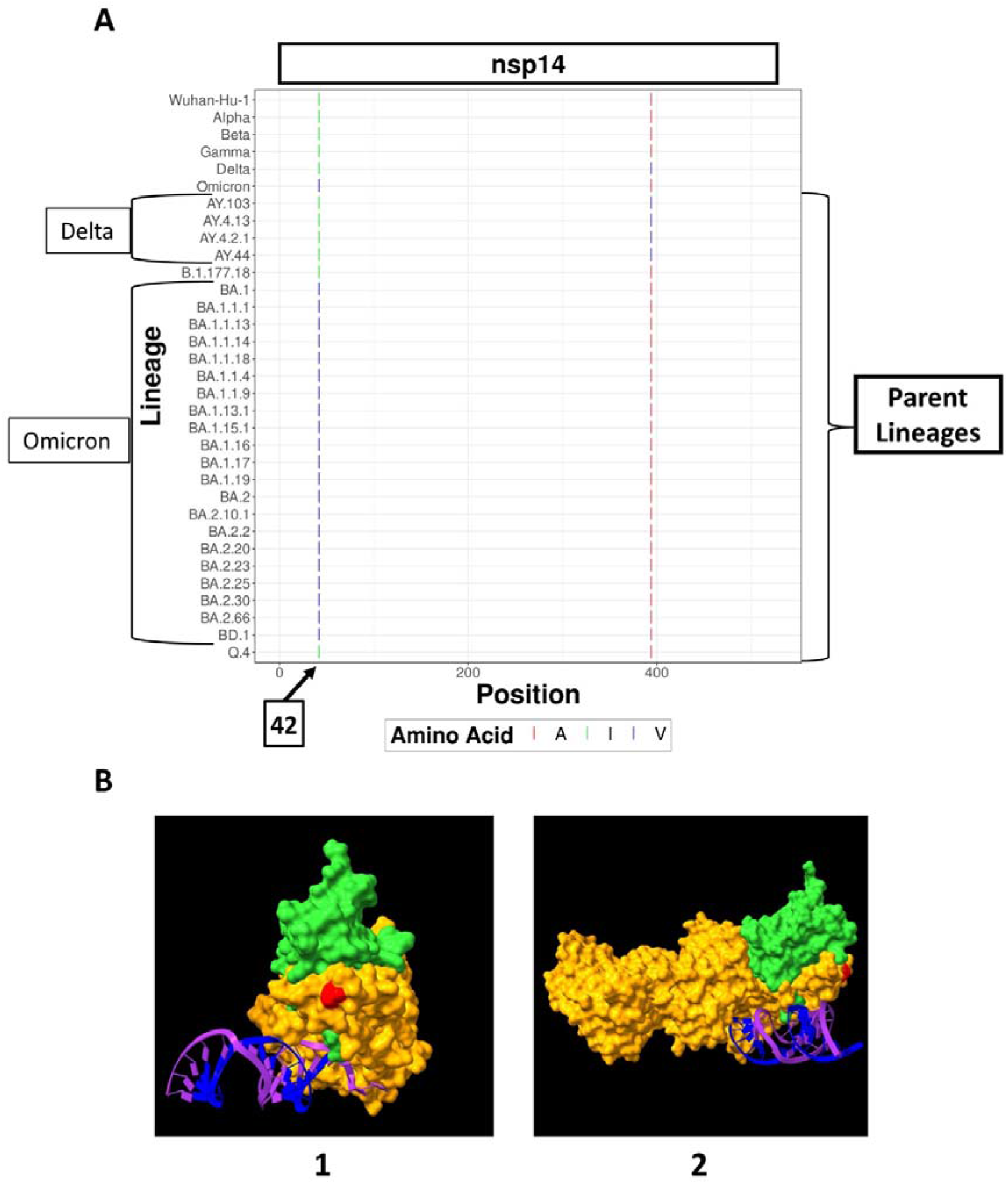
SARS-CoV2 Nsp14 recombinant parent lineage amino acid polymorphism compared to VOCs: (A) Recombinant parent Inter-lineage amino acid polymorphic positions in Nsp14 with respect to reference Wuhan-hu 1 strain sequence, Alpha Variant(B. 1.1.7), Beta Variant(B. 1.351), Gamma Variant(P. 1), Delta Variant(B.1.617.2) and Omicron Variant(B.1.1.529) were marked with “|”, with different colours representing different amino acids(Colour legends mark amino acids with one letter amino acid codes).X axis represent amion acid postion and Y axis represent lineages. Amino acid residue specifically conserved in omicron is named and pointed on the x-axis. (B) 3D structural visualisation of SARS-CoV2 Nsp10-Nsp14-RNA complex showing amino acid position conserved over all the omicron recombinant parent genomes. Here orange coloured residues represent Nsp14, lime-green-coloured residues represent Nsp10, magenta-coloured RNA strands represent Template strand(T-strand), blue-coloured RNA strands represent Product strand(P-strand) and red-coloured residues show the conserved residues in Nsp14. There are 2 insets: Inset 1 – Top view of Nsp10-Nsp14-RNA complex(PDB ID–7N0B); Inset 2 – Side view of Nsp10-Nsp14-RNA complex(PDB ID–7N0B).

**Supplementary Figure 5:**
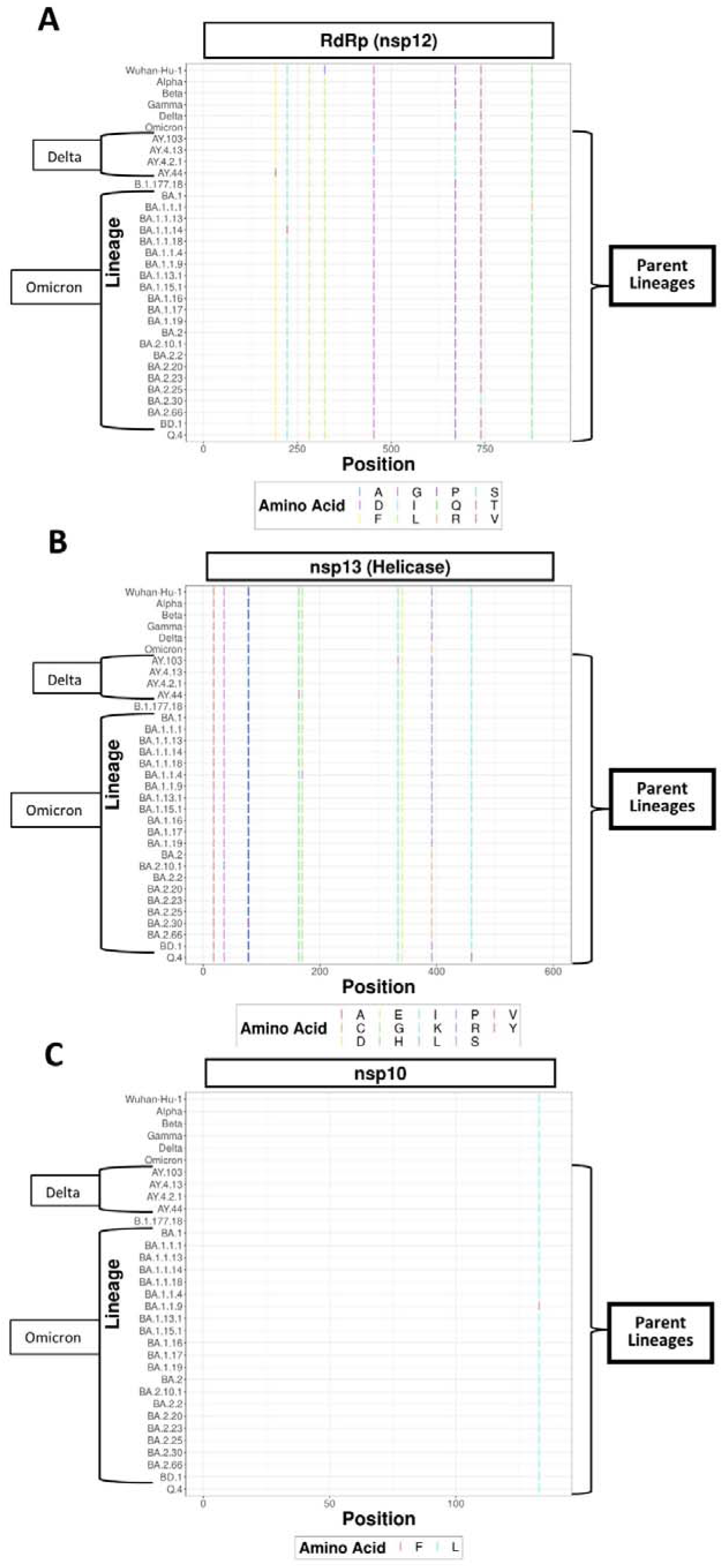
SARS-CoV2 RdRp, Helicase and Nsp10 recombinant parent lineage amino acid polymorphism compared to VOCs: Recombinant parent Inter-lineage amino acid polymorphic positions in the respective protein with respect to reference Wuhan-hu 1 strain sequence, Alpha Variant(B. 1.1.7), Beta Variant(B. 1.351), Gamma Variant(P.1), Delta Variant(B. 1.617.2) and Omicron Variant(B.1.1.529) were marked with “|”, with different colours representing different amino acids(Colour legends mark amino acids with one letter amino acid codes) X axis represent amino acid postion and Y axis represent lineages.. Legends are positioned to the bottom of each of the figures **(A)** Nsp12 (RdRp) **(B)** Nsp13 (Helicase) **(C)** Nsp 10

**Supplementary Table 1:**
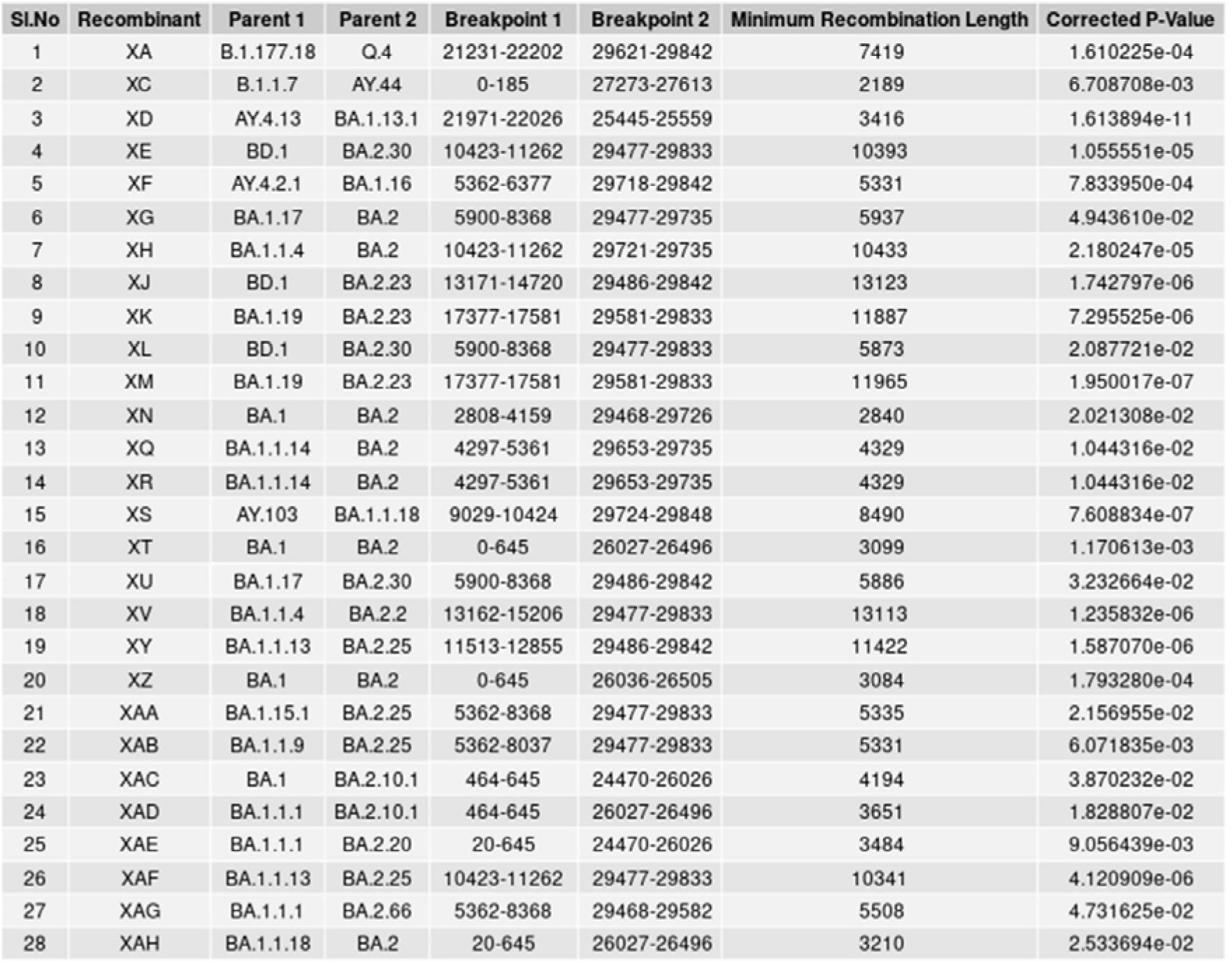
Identified best recombination parent lineages and breakpoint regions of each recombinant lineages inferred according to 3SEQ.Column 1: Serial number; Column 2: Recombinant lineage that results from recombination; Column 3: First Parent lineage involved in recombination; Column 4: Second Parent lineage involved in recombination; Column 5: Region of the first Breakpoint in the genome; Column 6: Region of the second Breakpoint in the genome; Column 7: Minimum length of the recombinant segments; Column 8: Dunn-Sidak correction of p, in mantissa-exponent format

| Sl.No | Recombinant | Parent 1 | Parent 2 | Breakpoint 1 | Breakpoint 2 | Minimum Recombination Length | Corrected P-Value |
| --- | --- | --- | --- | --- | --- | --- | --- |
| 1 | XA | B.1.177.18 | Q.4 | 21231-22202 | 29621-29842 | 7419 | 1.610225e-04 |
| 2 | XC | B.1.1.7 | AY.44 | 0-185 | 27273-27613 | 2189 | 6.708708e-03 |
| 3 | XD | AY.4.13 | BA.1.13.1 | 21971-22026 | 25445-25559 | 3416 | 1.613894e-11 |
| 4 | XE | BD.1 | BA.2.30 | 10423-11262 | 29477-29833 | 10393 | 1.055551e-05 |
| 5 | XF | AY.4.2.1 | BA.1.16 | 5362-6377 | 29718-29842 | 5331 | 7.833950e-04 |
| 6 | XG | BA.1.17 | BA.2 | 5900-8368 | 29477-29735 | 5937 | 4.943610e-02 |
| 7 | XH | BA.1.1.4 | BA.2 | 10423-11262 | 29721-29735 | 10433 | 2.180247e-05 |
| 8 | XJ | BD.1 | BA.2.23 | 13171-14720 | 29486-29842 | 13123 | 1.742797e-06 |
| 9 | XK | BA.1.19 | BA.2.23 | 17377-17581 | 29581-29833 | 11887 | 7.295525e-06 |
| 10 | XL | BD.1 | BA.2.30 | 5900-8368 | 29477-29833 | 5873 | 2.087721e-02 |
| 11 | XM | BA.1.19 | BA.2.23 | 17377-17581 | 29581-29833 | 11965 | 1.950017e-07 |
| 12 | XN | BA.1 | BA.2 | 2808-4159 | 29468-29726 | 2840 | 2.021308e-02 |
| 13 | XQ | BA.1.1.14 | BA.2 | 4297-5361 | 29653-29735 | 4329 | 1.044316e-02 |
| 14 | XR | BA.1.1.14 | BA.2 | 4297-5361 | 29653-29735 | 4329 | 1.044316e-02 |
| 15 | XS | AY.103 | BA.1.1.18 | 9029-10424 | 29724-29848 | 8490 | 7.608834e-07 |
| 16 | XT | BA.1 | BA.2 | 0-645 | 26027-26496 | 3099 | 1.170613e-03 |
| 17 | XU | BA.1.17 | BA.2.30 | 5900-8368 | 29486-29842 | 5886 | 3.232664e-02 |
| 18 | XV | BA.1.1.4 | BA.2.2 | 13162-15206 | 29477-29833 | 13113 | 1.235832e-06 |
| 19 | XY | BA.1.1.13 | BA.2.25 | 11513-12855 | 29486-29842 | 11422 | 1.587070e-06 |
| 20 | XZ | BA.1 | BA.2 | 0-645 | 26036-26505 | 3084 | 1.793280e-04 |
| 21 | XAA | BA.1.15.1 | BA.2.25 | 5362-8368 | 29477-29833 | 5335 | 2.156955e-02 |
| 22 | XAB | BA.1.1.9 | BA.2.25 | 5362-8037 | 29477-29833 | 5331 | 6.071835e-03 |
| 23 | XAC | BA.1 | BA.2.10.1 | 464-645 | 24470-26026 | 4194 | 3.870232e-02 |
| 24 | XAD | BA.1.1.1 | BA.2.10.1 | 464-645 | 26027-26496 | 3651 | 1.828807e-02 |
| 25 | XAE | BA.1.1.1 | BA.2.20 | 20-645 | 24470-26026 | 3484 | 9.056439e-03 |
| 26 | XAF | BA.1.1.13 | BA.2.25 | 10423-11262 | 29477-29833 | 10341 | 4.120909e-06 |
| 27 | XAG | BA.1.1.1 | BA.2.66 | 5362-8368 | 29468-29582 | 5508 | 4.731625e-02 |
| 28 | XAH | BA.1.1.18 | BA.2 | 20-645 | 26027-26496 | 3210 | 2.533694e-02 |

**Supplementary Table 2:**
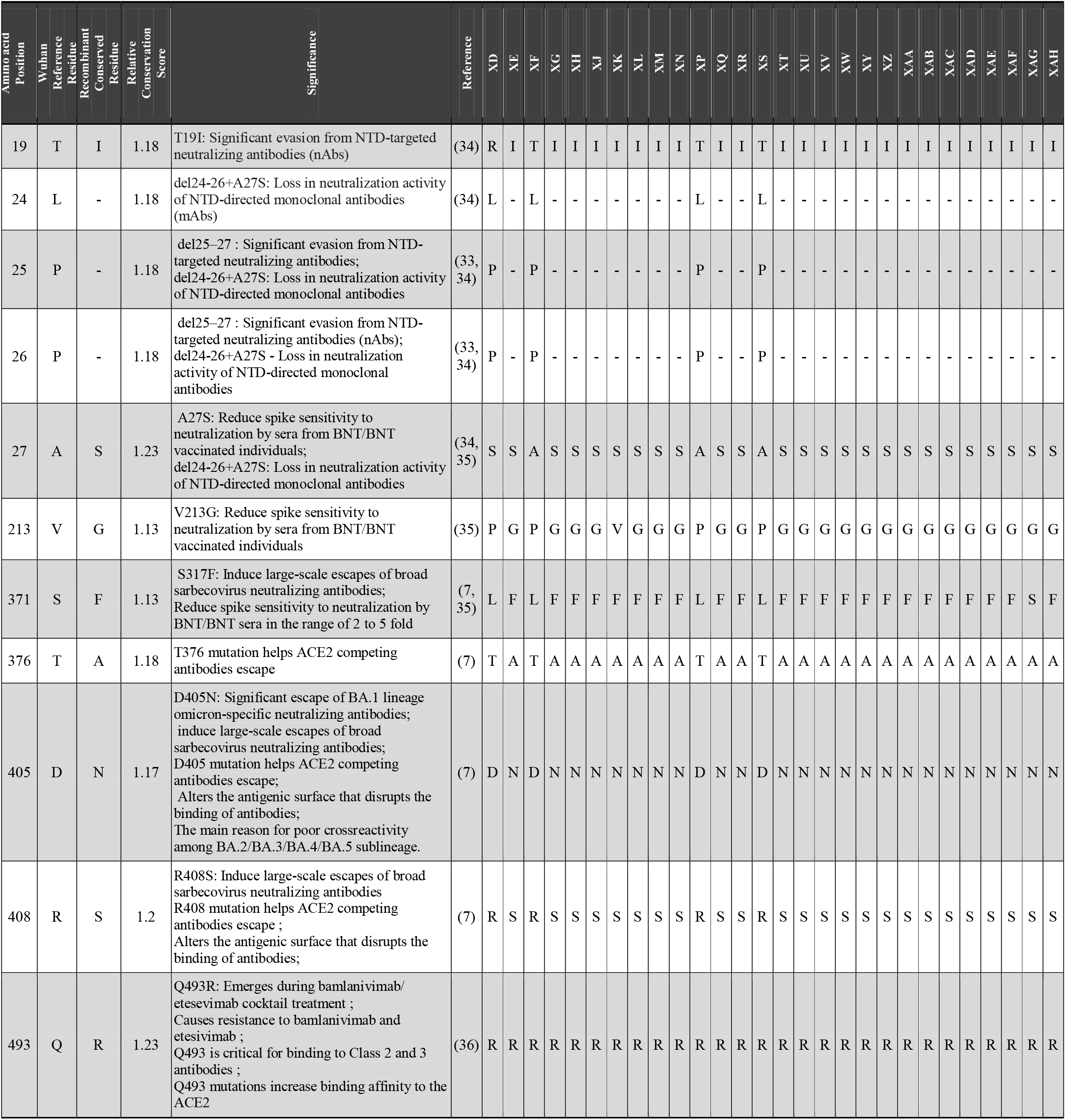
At least one omicron parenting Recombinant lineage spike residue conservation relative to omicron with discovered relevance in viral transmission and immune escape having recombinant specific residue information. Each conserved amino acid position in spike which is varying from Wuhan reference sequence wuhan reference spike residue at those positions in one letter code, conserved recombinant lineages residue at that position in one letter code, relative residue conservation score of recombinant lineages spike relative to residue conservation scores in spikes of omicron lineages, mutation significance, reference for the mutation significance information and specific amino acid residue in each of the 28 at least one omicron parenting recombinants in these conserved positions are tabulated.

