## Supplementary figures and images for "Enhanced Recombination Among SARS-CoV-2 Omicron Variants Contributes to Viral Immune Escape"

### S1

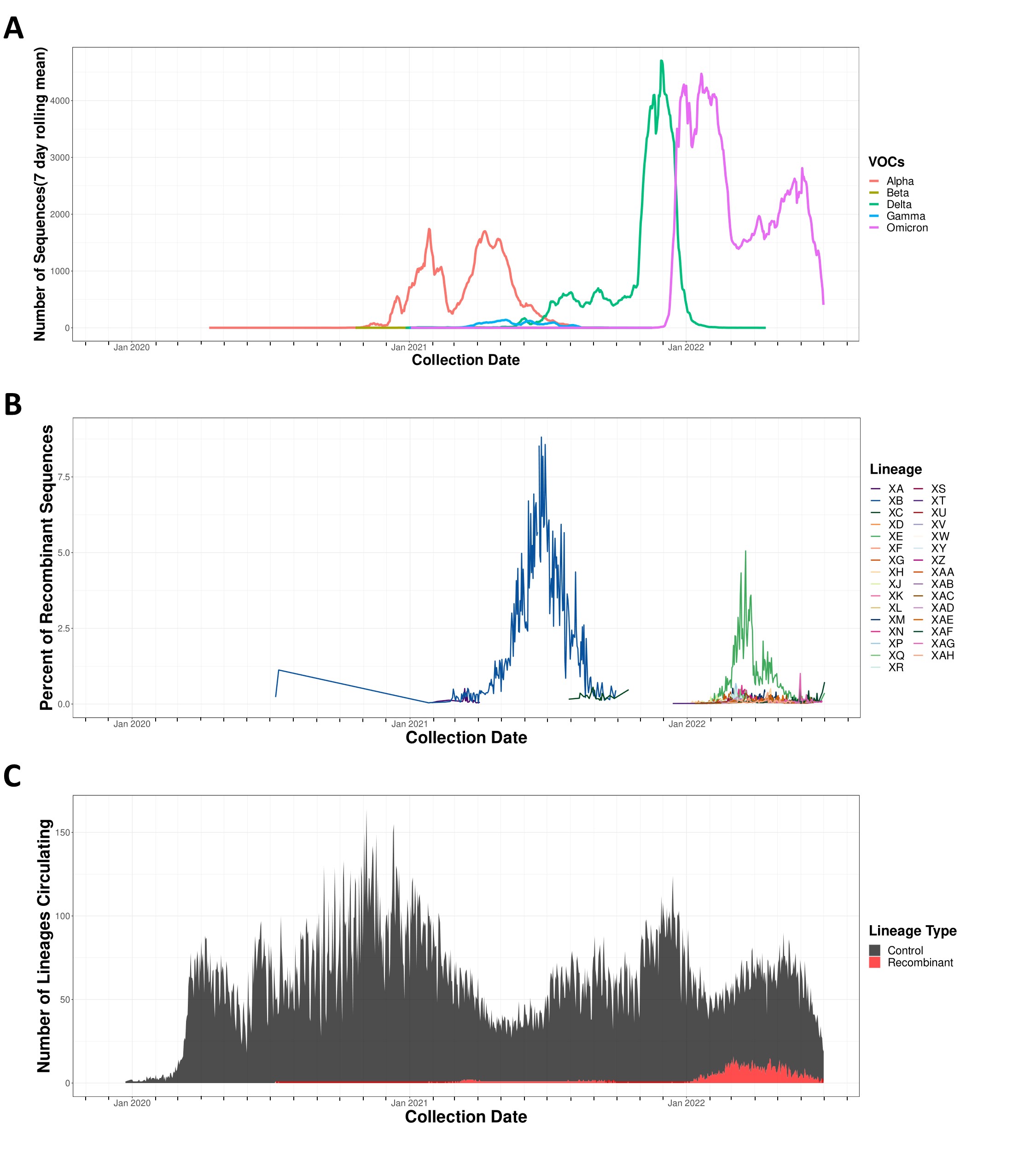

### S2A

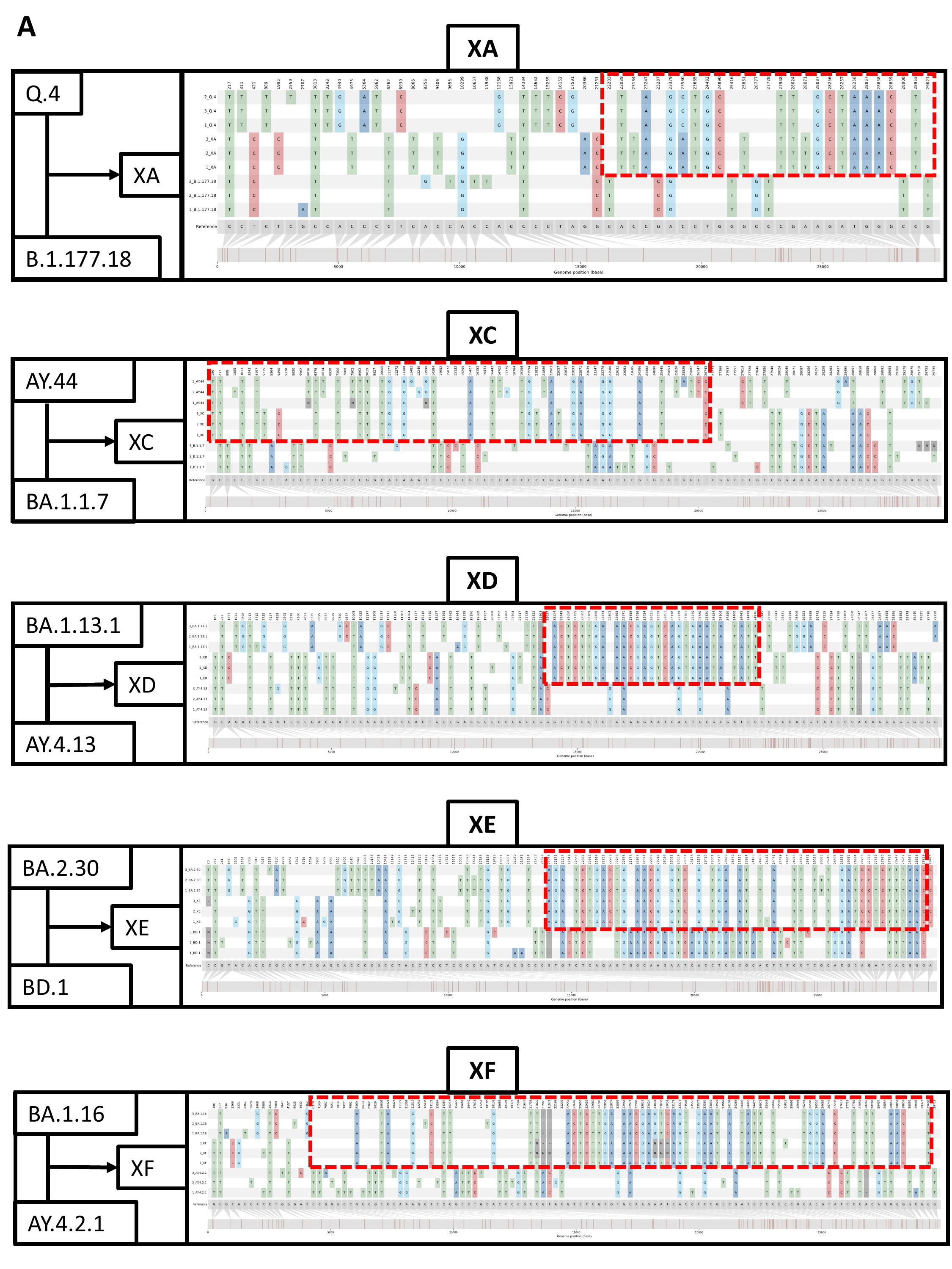

### S2B

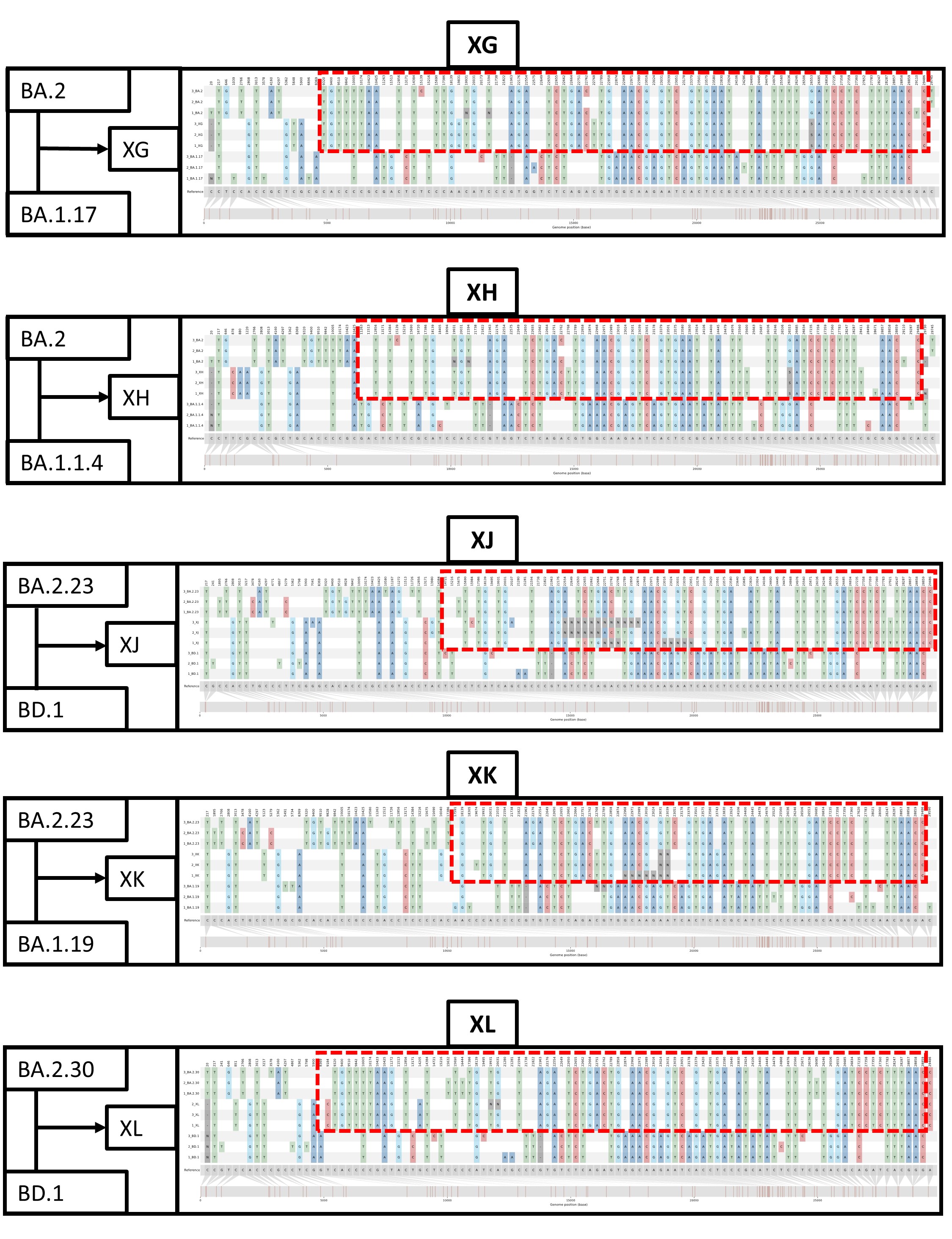

### S2C

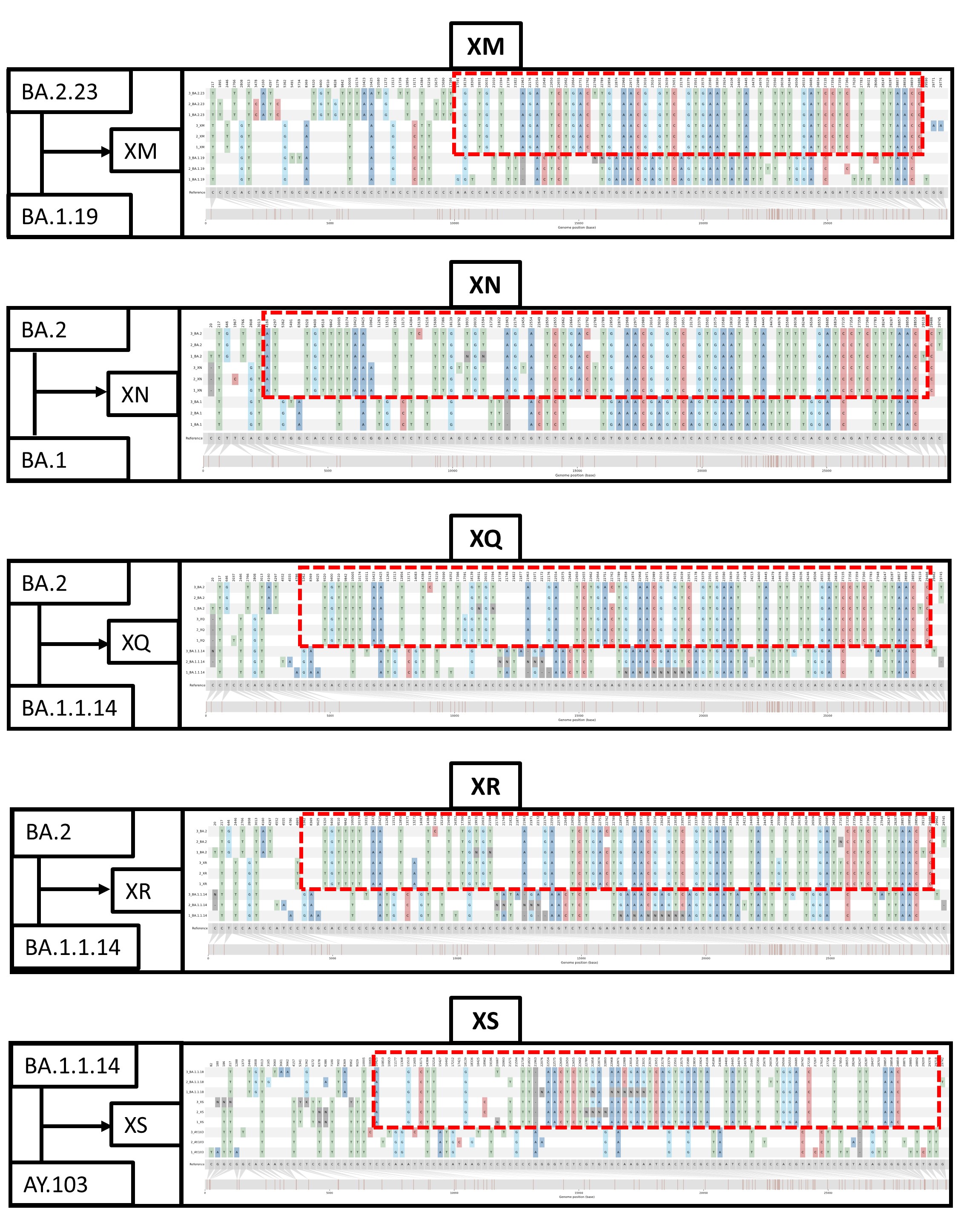

### S2D

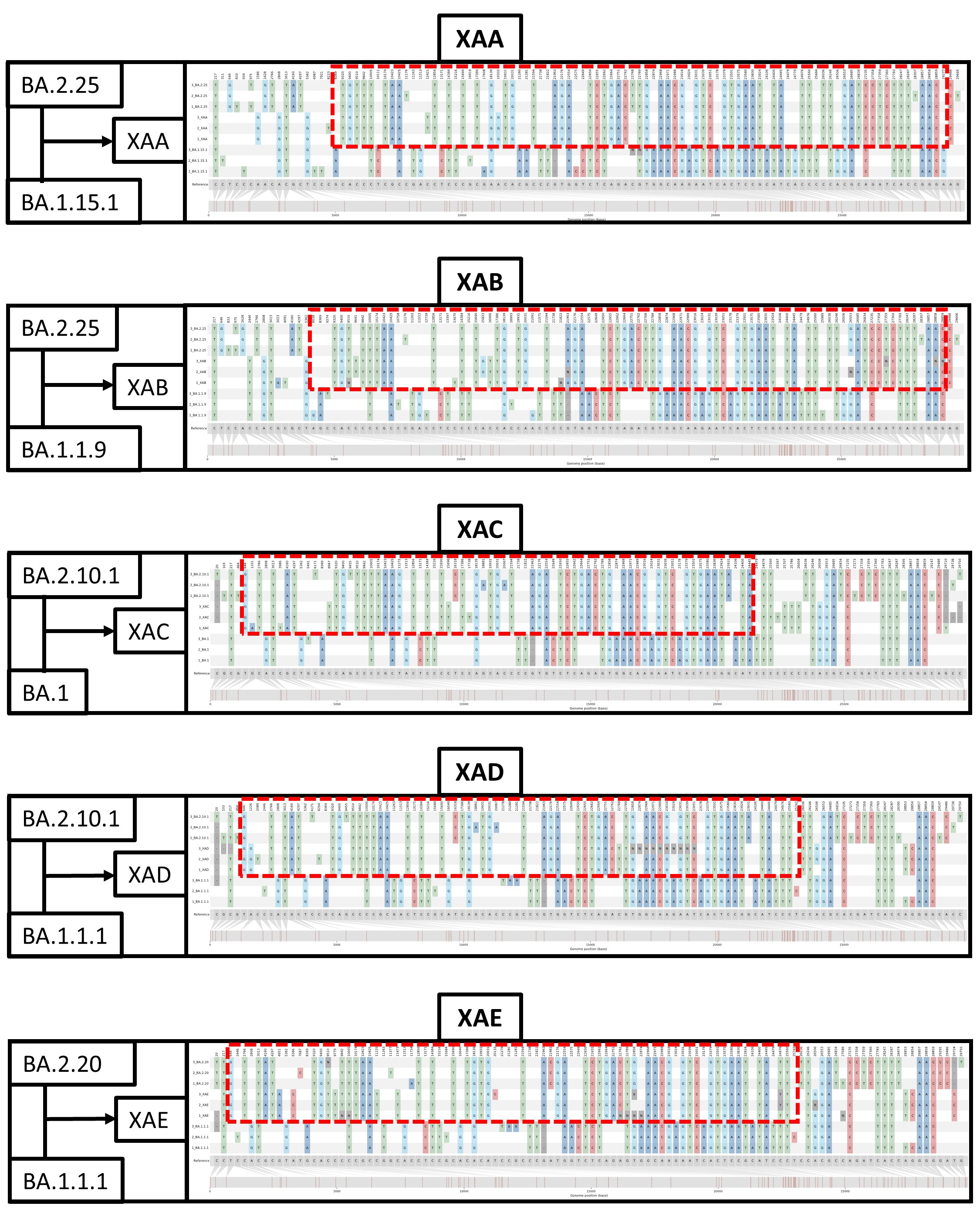

### S2E

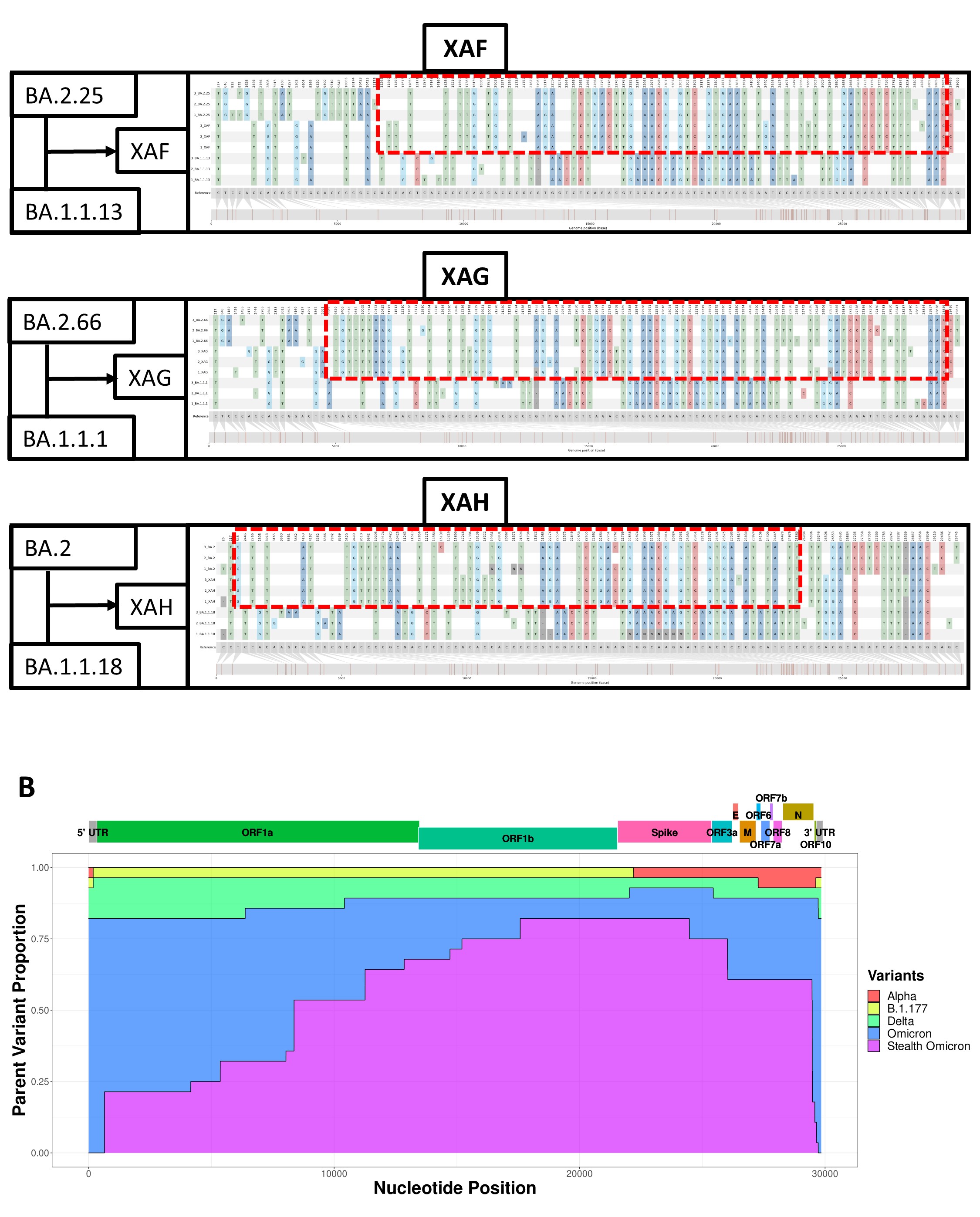

### S3

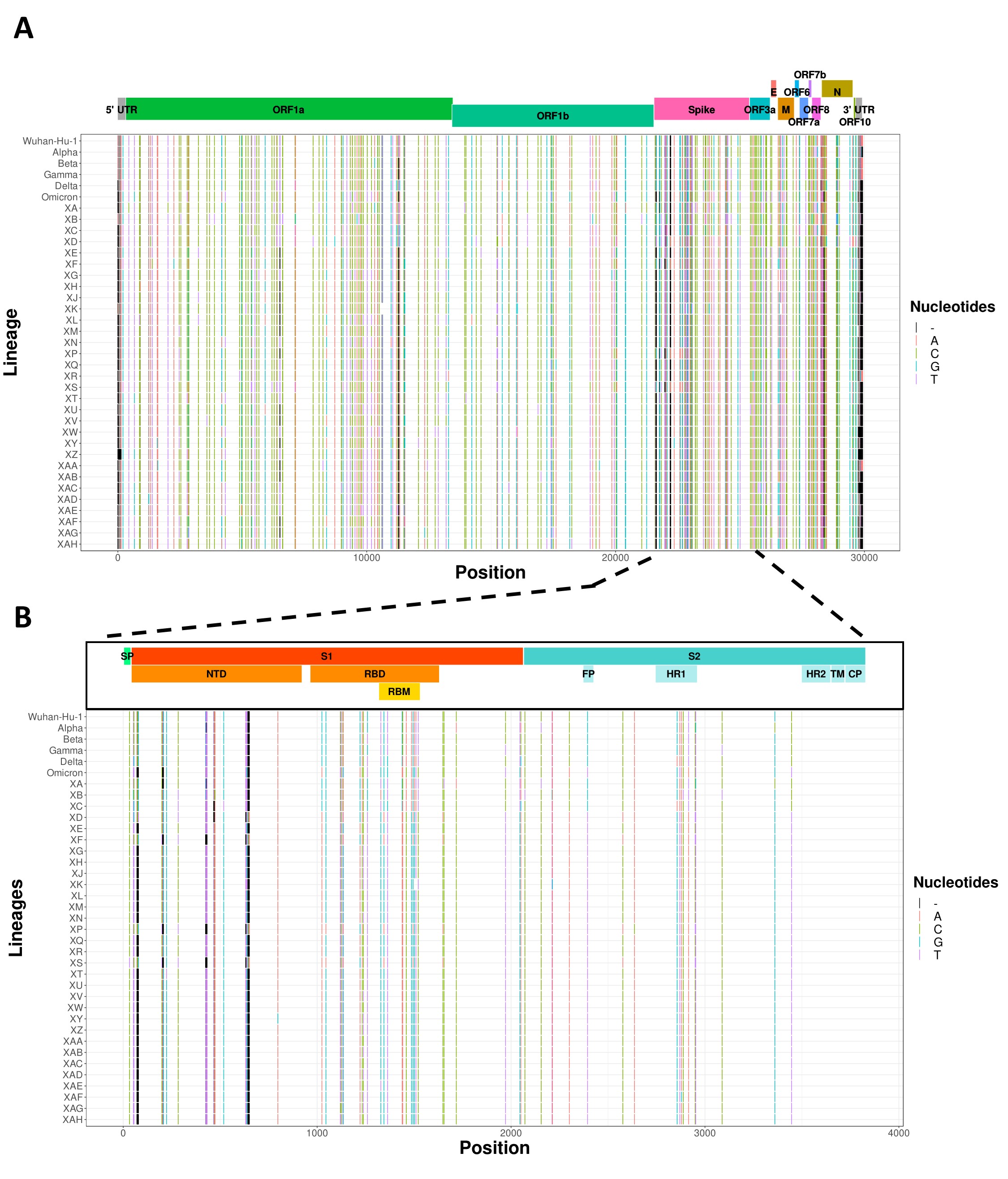

### S4

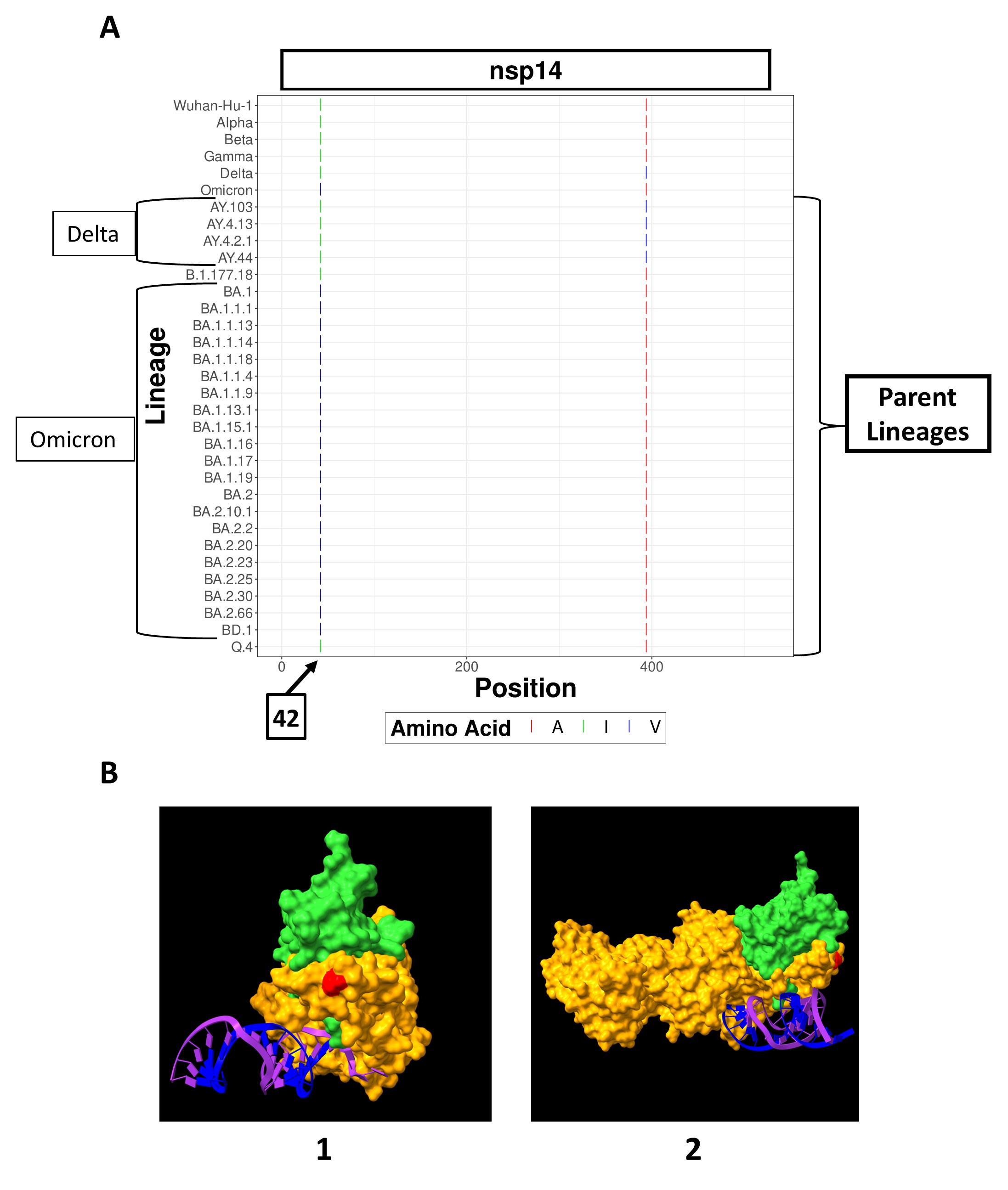

### S5

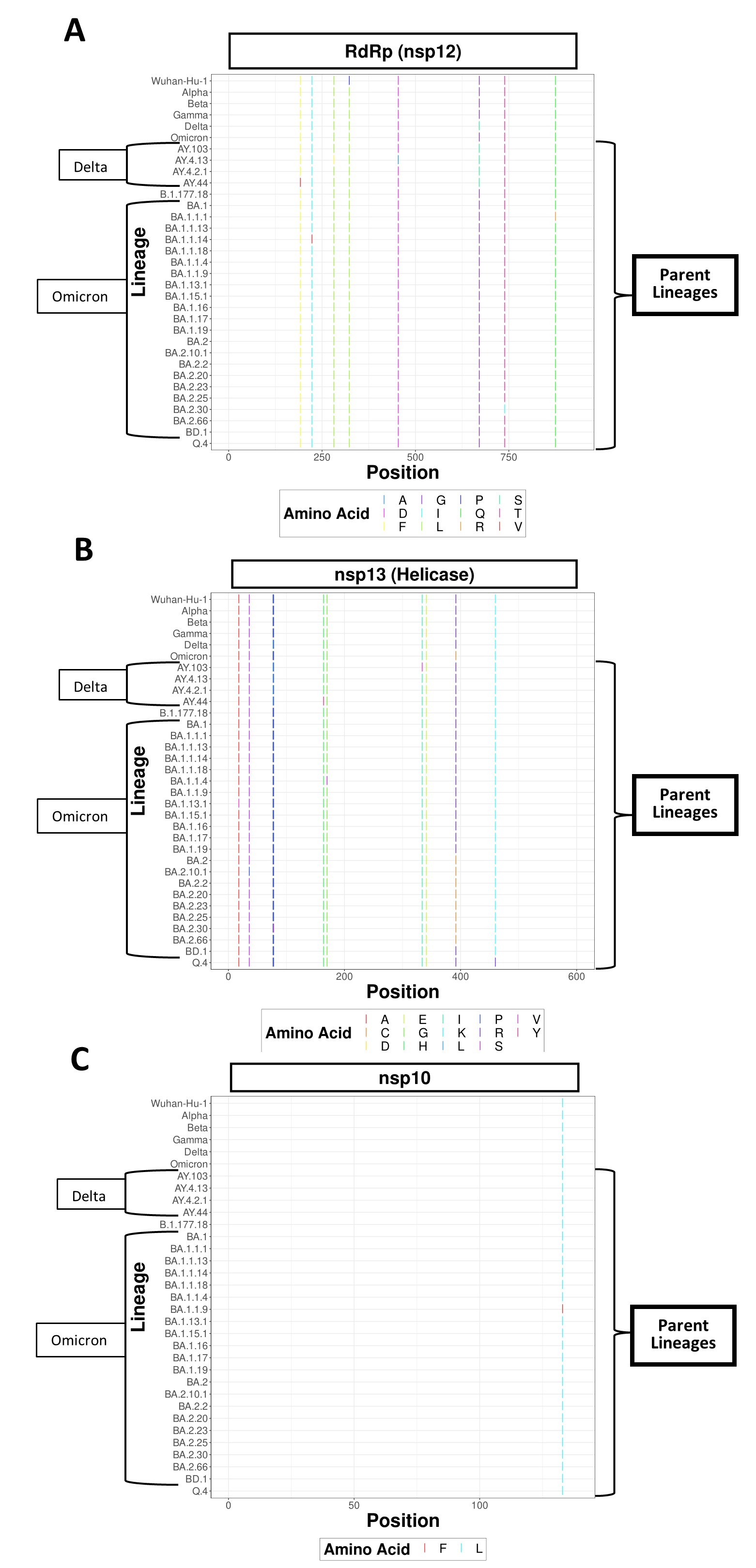

### Sup Table 1

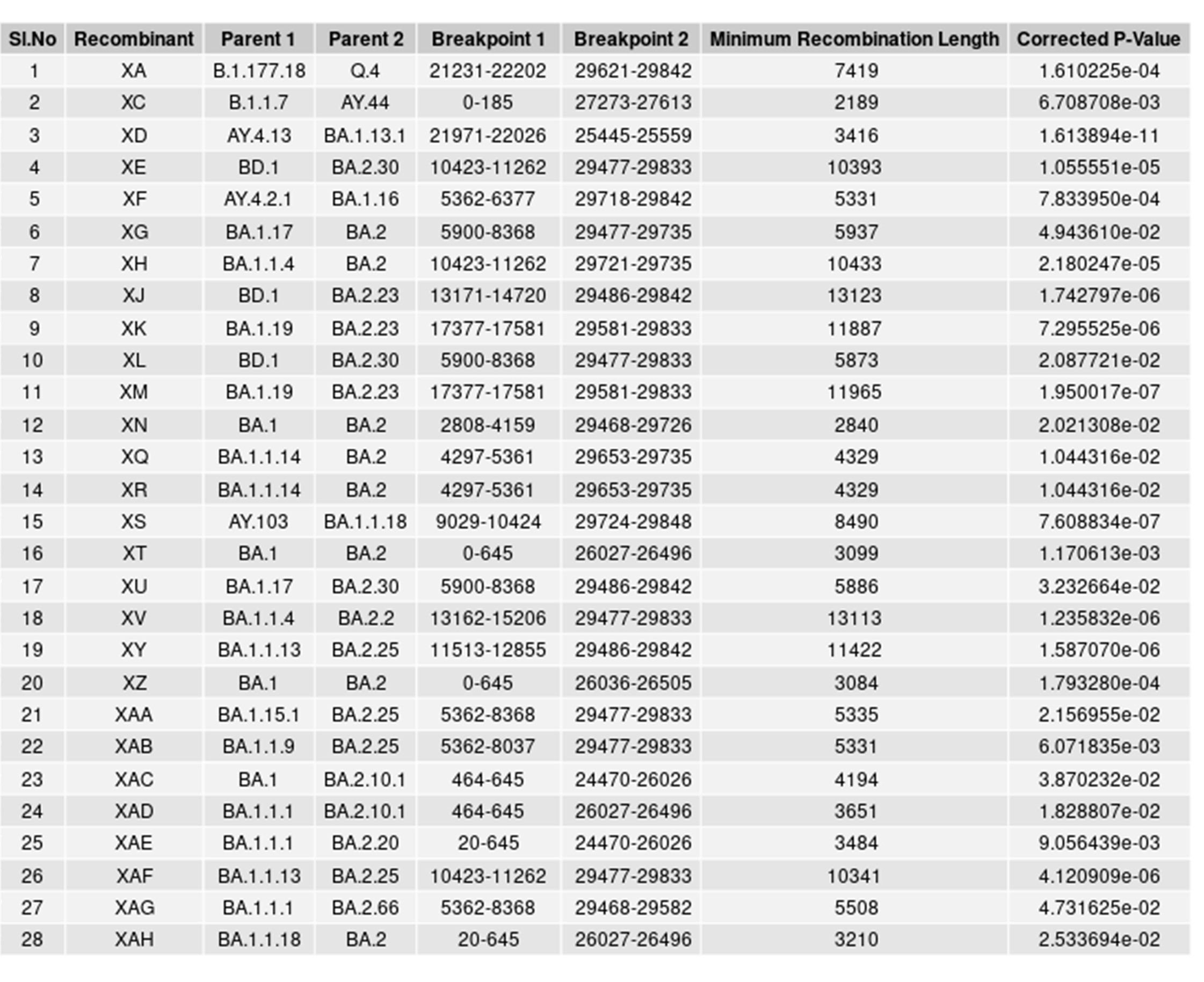
