## Supplementary material for "Enhanced Recombination Among SARS-CoV-2 Omicron Variants Contributes to Viral Immune Escape": Sup Table 2

| **Amino acid Position** | **Wuhan Reference Residue** | **Recombinant Conserved Residue** | **Relative Conservation  Score** | **Significance** | **Reference** | **XD** | **XE** | **XF** | **XG** | **XH** | **XJ** | **XK** | **XL** | **XM** | **XN** | **XP** | **XQ** | **XR** | **XS** | **XT** | **XU** | **XV** | **XW** | **XY** | **XZ** | **XAA** | **XAB** | **XAC** | **XAD** | **XAE** | **XAF** | **XAG** | **XAH** |
| --- | --- | --- | --- | --- | --- | --- | --- | --- | --- | --- | --- | --- | --- | --- | --- | --- | --- | --- | --- | --- | --- | --- | --- | --- | --- | --- | --- | --- | --- | --- | --- | --- | --- |
| 19 | T | I | 1.18 | T19I: Significant evasion from NTD-targeted neutralizing antibodies (nAbs) | (34) | R | I | T | I | I | I | I | I | I | I | T | I | I | T | I | I | I | I | I | I | I | I | I | I | I | I | I | I |
| 24 | L | - | 1.18 | del24-26+A27S: Loss in neutralization activity of NTD-directed monoclonal antibodies(mAbs) | (34) | L | - | L | - | - | - | - | - | - | - | L | - | - | L | - | - | - | - | - | - | - | - | - | - | - | - | - | - |
| 25 | P | - | 1.18 | del25–27 : Significant evasion from NTD-targeted neutralizing antibodies (nAbs) ;  del24-26+A27S: Loss in neutralization activity of NTD-directed monoclonal antibodies(mAbs) | (33, 34) | P | - | P | - | - | - | - | - | - | - | P | - | - | P | - | - | - | - | - | - | - | - | - | - | - | - | - | - |
| 26 | P | - | 1.18 | del25–27 : Significant evasion from NTD-targeted neutralizing antibodies (nAbs); del24-26+A27S - Loss in neutralization activity of NTD-directed monoclonal antibodies(mAbs) | (33, 34) | P | - | P | - | - | - | - | - | - | - | P | - | - | P | - | - | - | - | - | - | - | - | - | - | - | - | - | - |
| 27 | A | S | 1.23 | A27S: Reduce spike sensitivity to neutralization by sera from BNT/BNT vaccinated individuals; del24-26+A27S: Loss in neutralization activity of NTD-directed monoclonal antibodies(mAbs) | (34, 35) | S | S | A | S | S | S | S | S | S | S | A | S | S | A | S | S | S | S | S | S | S | S | S | S | S | S | S | S |
| 213 | V | G | 1.13 | V213G: Reduce spike sensitivity to neutralization by sera from BNT/BNT vaccinated individuals | (35) | P | G | P | G | G | G | V | G | G | G | P | G | G | P | G | G | G | G | G | G | G | G | G | G | G | G | G | G |
| 371 | S | F | 1.13 | S317F: Induce large-scale escapes of broad sarbecovirus neutralizing antibodies(nAbs) ; Reduce spike sensitivity to neutralization by BNT/BNT sera in the range of 2 to 5 fold | (7, 35) | L | F | L | F | F | F | F | F | F | F | L | F | F | L | F | F | F | F | F | F | F | F | F | F | F | F | S | F |
| 376 | T | A | 1.18 | T376 mutation helps ACE2 competing antibodies escape | (7) | T | A | T | A | A | A | A | A | A | A | T | A | A | T | A | A | A | A | A | A | A | A | A | A | A | A | A | A |
| 405 | D | N | 1.17 | D405N: Significant escape of BA.1 lineage omicron-specific neutralizing antibodies (nAbs) ;  induce large-scale escapes of broad sarbecovirus neutralizing antibodies(nAbs) ; D405 mutation helps ACE2 competing antibodies escape ;  Alters the antigenic surface that disrupts the binding of antibodies; The main reason for poor crossreactivity among BA.2/BA.3/BA.4/BA.5 sublineage. | (7) | D | N | D | N | N | N | N | N | N | N | D | N | N | D | N | N | N | N | N | N | N | N | N | N | N | N | N | N |
| 408 | R | S | 1.2 | R408S: Induce large-scale escapes of broad sarbecovirus neutralizing antibodies(nAbs) R408 mutation helps ACE2 competing antibodies escape ; Alters the antigenic surface that disrupts the binding of antibodies; | (7) | R | S | R | S | S | S | S | S | S | S | R | S | S | R | S | S | S | S | S | S | S | S | S | S | S | S | S | S |
| 493 | Q | R | 1.23 | Q493R: Emerges during bamlanivimab/etesevimab cocktail treatment ; Causes resistance to bamlanivimab and etesivimab ; Q493 is critical for binding to Class 2 and 3 antibodies ; Q493 mutations increase binding affinity to the ACE2 | (36) | R | R | R | R | R | R | R | R | R | R | R | R | R | R | R | R | R | R | R | R | R | R | R | R | R | R | R | R |

**Supplementary Table 2: At least one omicron parenting Recombinant lineage spike residue conservation relative to omicron with discovered relevance in viral transmission and immune escape having recombinant specific residue information**. Each conserved amino acid position in spike which is varying from Wuhan reference sequence wuhan reference spike residue at those positions in one letter code, conserved recombinant lineages residue at that position in one letter code, relative residue conservation score of recombinant lineages spike relative to residue conservation scores in spikes of omicron lineages, mutation significance, reference for the mutation significance information and specific amino acid residue in each of the 28 at least one omicron parenting recombinants in these conserved positions are tabulated.
